## Supplementary Data for "Activation by Cleavage of the Epithelial Na^+^ Channel α and γ Subunits Independently Coevolved with the Vertebrate Terrestrial Migration"

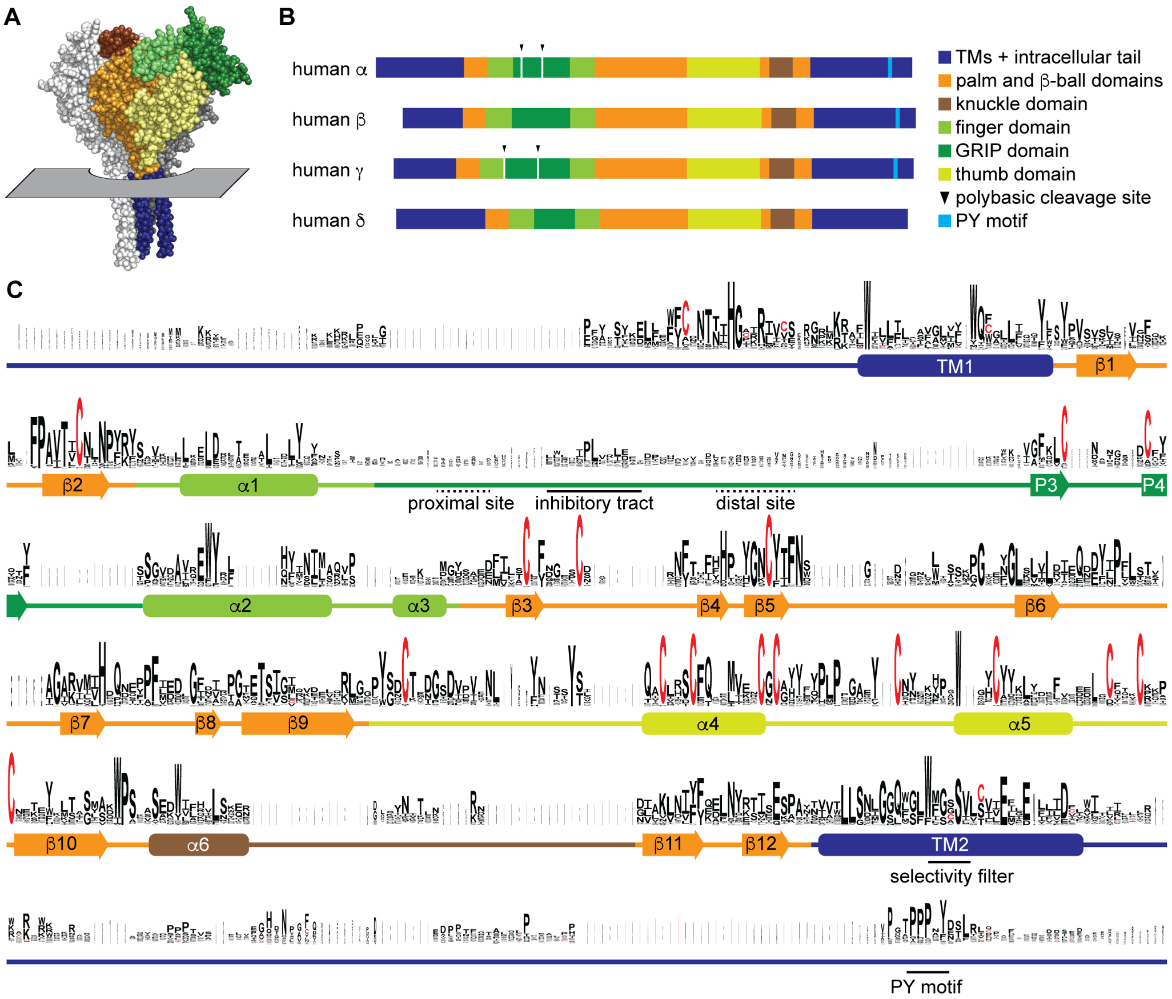


**Figure 1.** Sequence conservation in ENaC subunits. A, Space filling model of ENaC (pdb code: 6BQN) with plane indicating position of outer membrane border. The α and γ subunits are white and grey, respectively. The β subunit is colored by domain as indicated in panel B. Intracellular structures are absent in this structural model. B, Linear model of human ENaC subunits showing domain organization and highlighting position of polybasic cleavage sites and PY motifs. C, Sequences (Supplementary Table 1) were aligned using MUSCLE [21]. Residue symbol sizes are proportional to frequency at a given position. Key features in the sequence are indicated, as are the approximate position of α helices (rounded rectangles) and β-strands (arrows). Colors correspond to protein domains, as indicated in panel B. The GRIP domain is unique to ENaC subunits in the ENaC/Deg family. The P1 and P2 β-strands in the GRIP domain are not indicated, but are likely near the inhibitory tract and distal site, respectively, if present.

To examine the evolution of ENaC subunits, we assessed the phylogenetic relationship of our 53 assembled sequences with maximum-likelihood methods (Fig. 2). In the resulting tree, we observed 6 clades: ENaC $\alpha$, $\beta$, $\gamma$, and $\delta$ subunits, ASICs, and ENaC-like proteins. Each ENaC subunit clade largely recapitulated the expected phylogenetic relationships, with jawless fishes (lampreys) rooted closest to the base of the clade, followed by cartilaginous fishes (elephant shark), lobe-finned fishes (coelacanth and lungfishes), reptiles and birds, and mammals. Our evolutionary model suggests three gene duplication events for ENaC subunits, as previously reported [22]. The first duplication (node 1) gave ancestral $\beta/\gamma$ subunits (node 2) and $\alpha/\delta$ subunits (node 3). The second duplication (node 2) gave

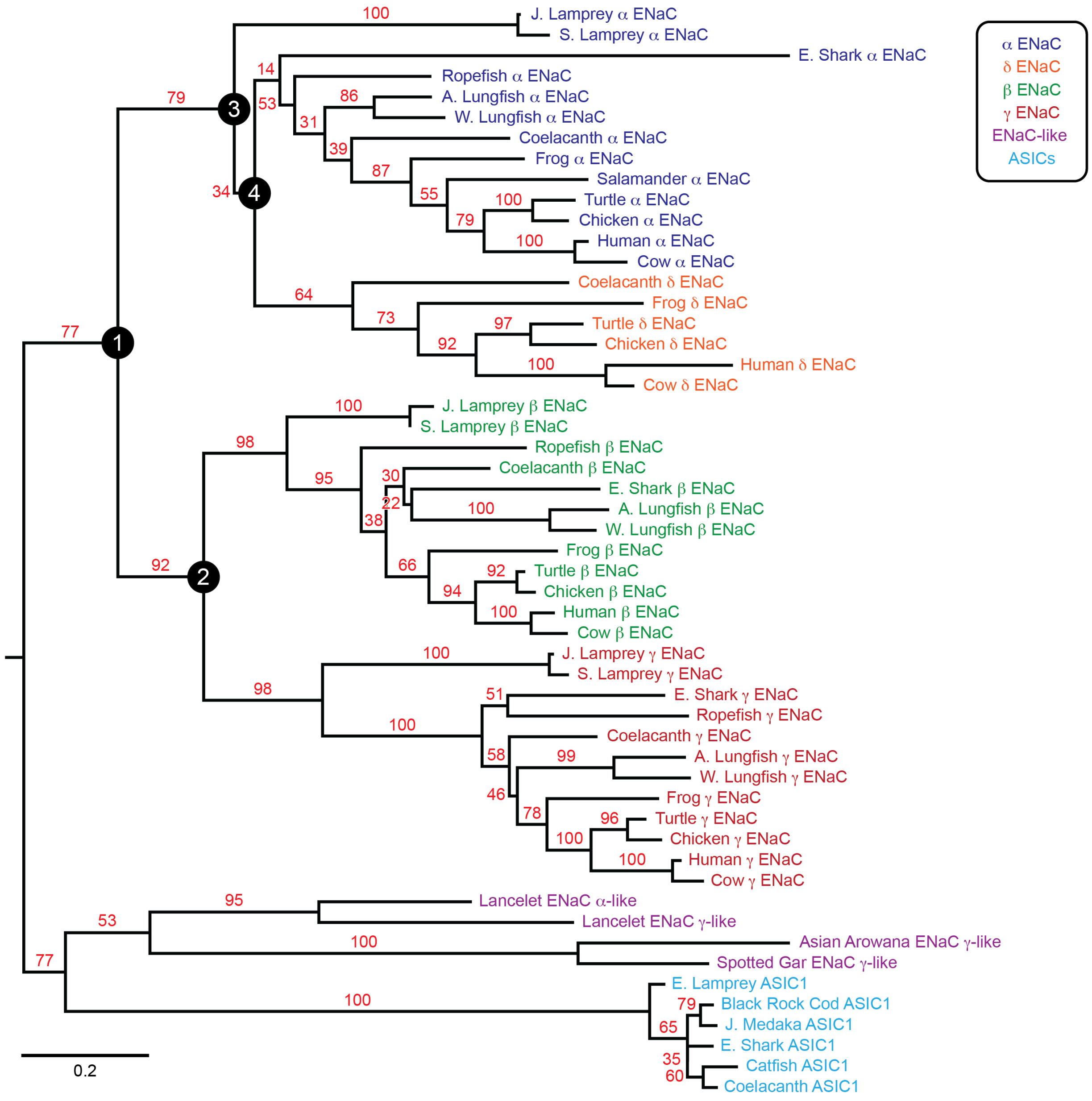


**Figure 2.** Phylogenetic tree of ENaC subunits. Maximum-likelihood tree calculated from ENaC subunit sequences of marine species and select terrestrial vertebrates, and ENaC-related proteins. Branch support bootstrap values are shown. Scale bar indicates the number of substitutions per site. Key ancestral nodes are indicated by circled numbers. A. Lungfish = Australian Lungfish, E. Shark = Elephant Shark, E. Lamprey = European River Lamprey, J. Lamprey = Japanese Lamprey, J. Medaka = Japanese Medaka, S. Lamprey = Sea Lamprey, W. Lungfish = West African Lungfish.

ancestral β and γ subunits. Both of these duplication events occurred in ancestors of jawless fishes at least 550 million years ago. The third duplication (node 4) leading to the divergence of α and δ subunits had more associated uncertainty. Given the presence of α and δ subunits in coelacanth, the duplication likely occurred before the divergence of the coelacanth and tetrapods.

*Polybasic tracts in the ENaC GRIP domains varied over time*

We then examined the sequence conservation of the polybasic tracts in the ENaC subunit GRIP domains required for ENaC activation by cleavage. In mammals, the GRIP domains of the α and γ subunits are each subject to double cleavage, leading to the release of embedded inhibitory tracts and channel activation. The β and δ subunits are not similarly processed. The proprotein convertase furin cleaves both the proximal and distal sites of the α subunit, and the proximal site of the γ subunit [7]. Cleavage distal to the γ subunit inhibitory tract can be catalyzed by several proteases at the cell surface, including prostasin at a polybasic tract [23, 24]. Similar results were reported for ENaC from *Xenopus laevis* [25]. We inspected our multiple sequence alignment for polybasic tracts that aligned closely with the tracts in mammalian and frog α and γ subunits (Supplementary Fig. 1). We found polybasic tracts aligning with the proximal and distal sites of the human α and γ subunits for all tetrapod α and γ subunits (Table 1). We also found both sites present in the γ subunit from *Erpetoichthys calabaricus* (Ropefish) and *Neoceratodus forsteri* (Australian lungfish), but not in *Protopterus annectens* (West African lungfish). In coelacanth, we identified single polybasic tracts in the α, β, and γ subunits. The elephant shark's γ subunit also had a single distal polybasic tract. The human δ subunit sequence also exhibits a polybasic tract in this region, but experimental evidence shows that it is not cleaved [26].

| | | | α | | δ | | β | | γ | |
|---|---|---|---|---|---|---|---|---|---|---|
| Animal | Terr. | Lungs | site 1 | site 2 | site 1 | site 2 | site 1 | site 2 | site 1 | site 2 |
| S. Lamprey | | | — | — | Ø | Ø | — | — | — | — |
| J. Lamprey | | | — | — | Ø | Ø | — | — | — | — |
| E. Shark | | | (···) | (···) | Ø | Ø | — | — | — | RQHR |
| Ropefish | | X | — | — | Ø | Ø | — | — | RKRR | NRKR |
| Coelacanth | | | RSNR | — | — | — | KRER | — | — | VKQR |
| W. Lungfish | | X | — | — | Ø | Ø | — | — | — | — |
| A. Lungfish | | X | — | — | Ø | Ø | — | — | RKLR | RQYR |
| Frog | X | X | RVKR | RVSR | — | — | — | — | RSKR | KRTR |
| Salamander | X | X | RERR | RVRR | Ø | Ø | Ø | Ø | Ø | Ø |
| Turtle | X | X | RSPR | RHKR | — | — | — | — | KVRR | NKRK |
| Chicken | X | X | RTSR | RQKR | — | — | — | — | KVRR | RKRK |
| Cow | X | X | RSRR | RGVR | — | *RLQR*[1] | — | — | RKRR | RKRK |
| Human | X | X | RSRR | RRAR | — | *RLQR*[1] | — | — | RKRR | RKRK |

**Table 1.** Polybasic sequences aligning with human ENaC subunit proximal (site 1) and distal (site 2) cleavage sites. Terr., terrestrial. Dashes indicate absence of polybasic tract. [1]The ENaC δ subunit in mammals has a polybasic sequence in the aligned region, but is not cleaved in human channels [26]. Ø, no sequence was available. (···), GRIP domain sequence was missing in the available sequence. Species abbreviations are as in Fig. 2.

*Australian lungfish ENaC γ subunit is cleaved at GRIP domain polybasic tracts*

To determine whether apparent cleavage sites are functional in a subunit rooted before the emergence of tetrapods, we examined the ENaC γ subunit from Australian lungfish (Aγ) in *Xenopus laevis* oocytes. Aγ has two furin cleavage motifs predicted to lead to activation (Table 1, Fig. 3A). To isolate functional effects to Aγ, we coexpressed an α subunit from mouse lacking residues excised by furin (mouse αΔ206-231; $m\alpha^{\Delta}$), rendering it incapable of proteolytic activation [27]. We also coexpressed a mouse β subunit truncated before the C-terminal PY motif (mouse βR564X; $m\beta^{T}$) to decrease channel turnover [28]. All γ subunits contained a C-terminal hemagglutinin (HA) epitope tag to facilitate detection (Fig. 3A). Oocytes expressing ENaC were conjugated with a membrane impermeant biotin reagent to

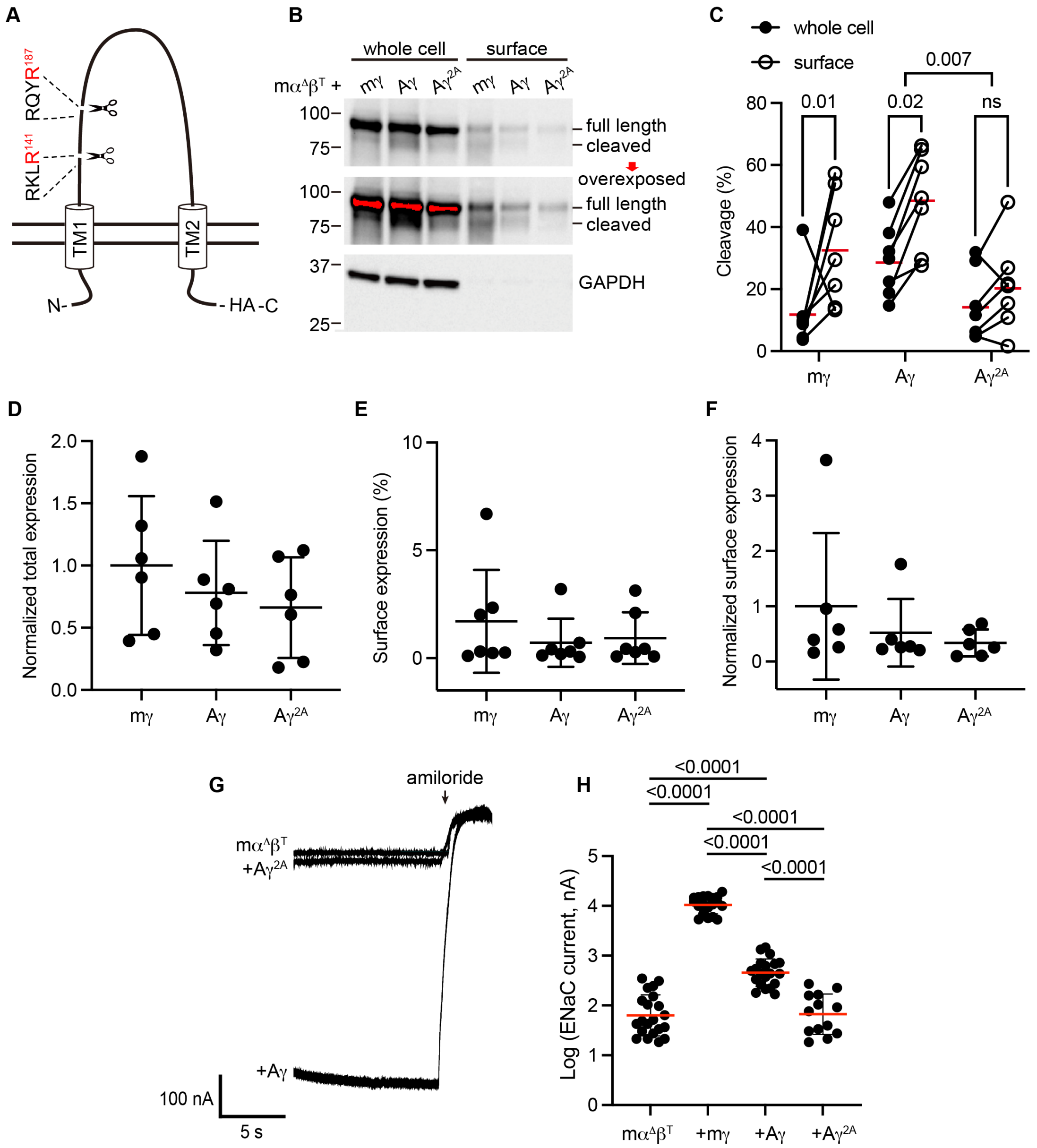


**Figure 3.** Predicted cleavage sites in the ENaC γ subunit from Australian lungfish are functional. A, Schematic of Aγ topology. Aγ has 2 predicted furin cleavage sites in its extracellular GRIP domain. All γ subunits were labeled with C-terminal epitope tags to facilitate detection of full-length subunits and the larger of the cleaved fragments. B, *Xenopus* oocytes were injected with cRNAs encoding $m\alpha^{\Delta}$, $m\beta^{T}$, and HA-tagged γ subunits, as indicated. One day after injection, whole cell lysates and cell surface isolates were blotted and probed for HA and GAPDH. Full length and cleaved bands are indicated and band densities were quantified. An overexposed blot is shown to highlight cell surface bands. Over exposed areas are red. C, Cleavage %, calculated as cleaved/(cleaved +full length)*100 is shown. Data were analyzed by repeated measures two-way ANOVA with Šidák's multiple comparison test. P-values are shown for indicated comparisons. Cleavage was also greater for Aγ than for mγ (p=0.05). D, Normalized total expression was calculated by normalizing the sum of full length and cleaved bands to the mean of mγ after normalizing each sample for loading based on GAPDH from the same blot. E, Surface expression % was calculated using band densities adjusted for the fraction of the respective sample loaded. F, Normalized surface expression was calculated by multiplying values form the same sample in D and E, and then normalizing to the mean of mγ. Data in D–F were analyzed by one-way ANOVA with Tukey's multiple comparison test. No significant differences between groups were found. Note that due to the lack of GAPDH data for one blot, the number of replicates for D and F (n=6) are one fewer than for C and E (n=7). G, Whole cell currents were measured in injected oocytes by two-electrode voltage clamp, with voltage clamped at -100 mV. Representative traces of indicated subunit combinations are shown. Currents were continuously recorded in a bath solution containing 110 mM $Na^{+}$. The ENaC-blocking drug amiloride (100 μM) was added at the end of each experiment to determine the ENaC-mediated current. H, Log transformed amiloride-sensitive inward currents are plotted, and were analyzed by one-way ANOVA followed by Tukey's multiple comparison test. P-values for comparisons where $p < 0.05$ are shown. Bars indicate mean values; errors shown are SD.

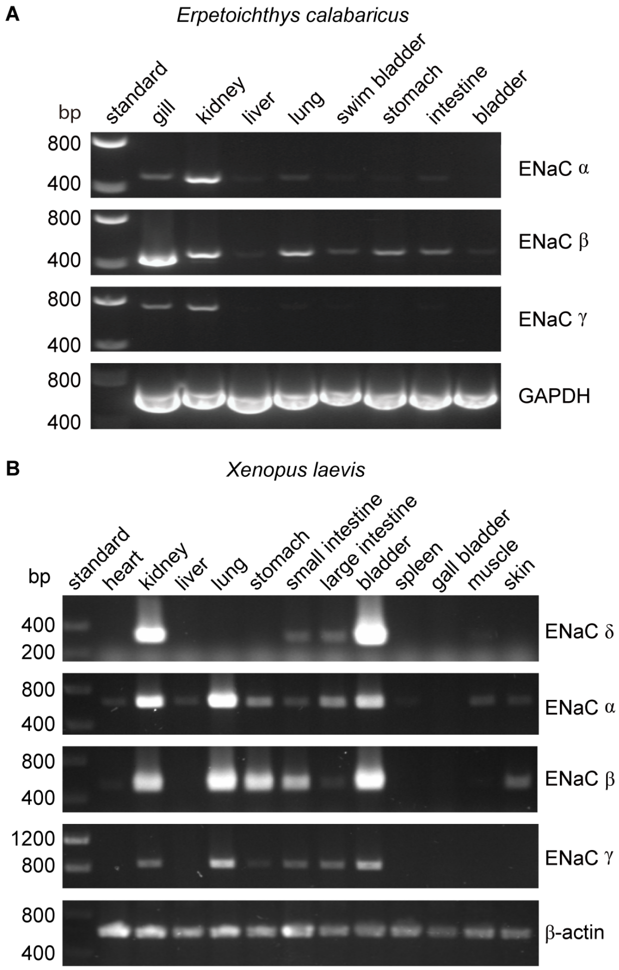


**Figure 4.** Tissue distribution of *Erpetoichthys calabaricus* (ropefish) and *Xenopus laevis* ENaC subunit transcripts by RT-PCR. cDNA libraries were generated from tissue homogenates. PCR reactions were performed using primers indicated in Supplementary Table 3.

| Animal | Terrestrial | Lungs | α | δ | β | γ |
|---|---|---|---|---|---|---|
| S. Lamprey | | | PPPSF | Ø | PPPHY | PPPQY |
| J. Lamprey | | | PPDY | Ø | PPPHY | PPPQY |
| E. Shark | | | (···) | Ø | PPPRY | PPPNY |
| Ropefish | | X | PPPAY | Ø | PPPHY | PPPNY |
| Coelacanth | | | PPAY | (···) | PPPNY | PPPTY |
| W. Lungfish | | X | PPPAY | Ø | PPPHY | PPPQY |
| A. Lungfish | | X | PPPAY | Ø | PPPKY | PPPQY |
| Frog | X | X | PPPAY | — | PPPNY | PPPKY |
| Salamander | X | X | PPPAY | Ø | Ø | Ø |
| Turtle | X | X | LPSY | — | PPPNY | PPPNY |
| Chicken | X | X | LPSY | — | PPPNY | PPPNY |
| Cow | X | X | PPPAY | — | PPPNY | PPPRY |
| Human | X | X | PPPAY | — | PPPNY | PPPKY |

**Table 2.** PY motifs (L/P-P-X-Y) in the C-terminal tails of ENaC subunits from various species. Ø, no sequence was available. (···), sequence for the C-terminal region was missing in the available sequence. Species abbreviations are as in Fig. 2.

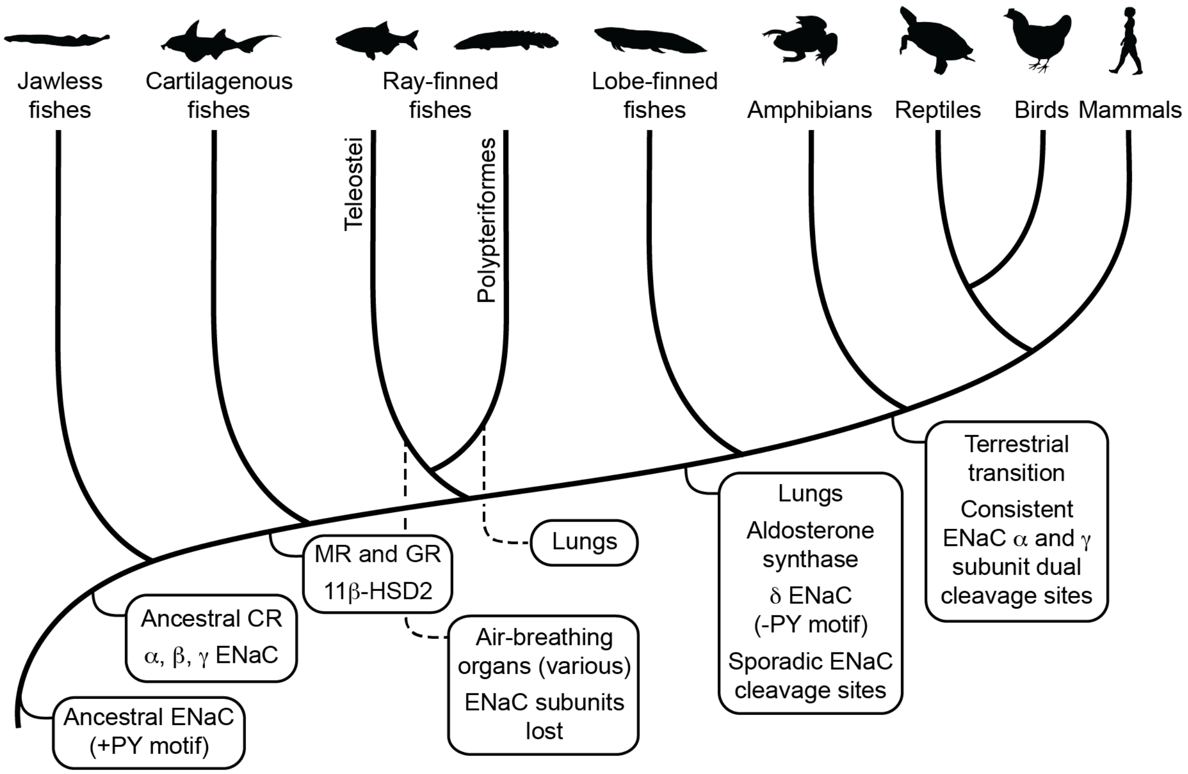


**Figure 5.** Schematic view of evolution of aldosterone signaling, air-breathing organs, ENaC, and ENaC regulatory motifs. ENaC subunits were not found in non-vertebrate chordates or in teleosts. The ancestral ENaC subunit likely had a PY motif and was a substrate for Nedd4-2-dependent regulation. Like mammalian ENaC α subunits, the ancient ENaC subunit may have formed functional homotrimers, or may have formed channels with other ENaC paralogs. ENaC α, β and γ subunits appeared before the emergence of jawless fishes, whereas ENaC δ subunits first appeared in an ancestor of the lobe-finned coelacanth. Proteins required for aldosterone signaling (MR, 11β-HSD2, and aldosterone synthase) evolved before the emergence of tetrapods. Lungs appeared in lobe-finned fishes on the lineage to tetrapods. Air-breathing organs (e.g. respiratory gas bladders and labyrinth organs) evolved independently in ray-finned fishes, including morphologically distinct lungs in Polypteriformes. Individual GRIP domain ENaC cleavage sites first appeared sporadically in marine species. Dual cleavage sites appeared consistently in the ENaC α and γ subunits in terrestrial vertebrates. CR, corticoid receptor; MR, mineralocorticoid receptor; GR, glucocorticoid receptor. Animal silhouettes courtesy of PhyloPic (http://www.phylopic.org).

*Motif analysis*

Furin cleaves human α ENaC after $RSRR^{178}$ and $RRAR^{204}$, and human γ ENaC at $RKRR^{138}$ [7]. Furin requires Arg at P1 and has a preferred P4-P3-P2-P1↓ = R-X-R/K-R↓ substrate sequence, although deviations at P2 and P4 have been observed, e.g. Ala-203 at P2 in the ENaC α subunit [68, 69]. Prostasin cleaves the human ENaC γ subunit after $RKRK^{181}$ [8]. Prostasin cleavage requires Arg or Lys at P1, and prefers basic residues at P2, P3, and P4 [60]. We inspected the region in the alignment that contains the human ENaC cleavage sites (Supplementary Fig. 1) for ideal furin sites or alternatively, for polybasic tracts ending in Arg that aligned within five residues of either of the $\alpha$ subunit furin sites for $\alpha$ and $\delta$ subunits, the $\gamma$ subunit furin site for $\beta$ and $\gamma$ subunits, or for any of the furin sites for ASIC subunits. We also inspected the alignment for polybasic tracts ending in Arg or Lys that aligned within five residues of the human $\gamma$ subunit prostasin site for $\beta$ and $\gamma$ subunits. For the PY motif (L/P-P-X-Y) required for Nedd4-2 dependent regulation [70], we inspected the C-terminal region of the alignment. All sequences found aligned with human ENaC subunit PY motifs in the C-termini.

*Plasmids and site-direct mutagenesis*

cDNAs encoding mouse ENaC subunits were previously described [28, 71, 72]. Australian lungfish $\gamma$ subunit with a C-terminal epitope tag (A$\gamma$) was synthesized by Twist Bioscience (San Francisco, CA) and cloned into pcDNA 3.1 hygro (+). Site-directed mutagenesis of A$\gamma$ was performed using the QuikChange


**Competing Interests**

The authors declare no conflicts of interest.

**References**


1. Long JA, Gordon MS. The greatest step in vertebrate history: a paleobiological review of the fish-tetrapod transition. Physiol Biochem Zool. 2004;77(5):700-19. Epub 2004/11/18. doi: 10.1086/425183. PubMed PMID: 15547790.

**Supplementary Table 1.** Protein sequences were found using BLAST tools at NCBI (https://www.ncbi.nlm.nih.gov/) and UniProt (https://www.uniprot.org/). [#]Annotated as ENaC $\alpha$ subunit in NCBI, but named ASIC1 here on the basis of calculated phylogenetic tree in Fig. 2. *Protein sequences were originally found using the BLAST tool at A*STAR (http://jlampreygenome.imcb.a-star.edu.sg/ and http://esharkgenome.imcb.a-star.edu.sg/), but were no longer available at the time of publication. Coding sequences are available at the accession numbers shown at NCBI.

| Species Name | Common Name | Protein | Accession Number | Database |
|---|---|---|---|---|
| *Callorhinchus milii* | Elephant Shark | ASIC1 | XP_007884967.1 | NCBI |
| *Lampetra fluviatilis* | European River Lamprey | ASIC1 | AAY28983.1 | NCBI |
| *Latimeria chalumnae* | Coelacanth | ASIC1 | XP_006007803.1 | NCBI |
| *Notothenia coriiceps* | Black Rock Cod | ASIC1 | XP_010778727.1 | NCBI |
| *Oryzias latipes* | Japanese Medaka | ASIC1 | XP_004068864.1 | NCBI |
| *Ictalurus punctatus* | Catfish | ASIC1[#] | XP_017320908.1 | NCBI |
| *Bos taurus* | Cow | ENaC $\alpha$ | NP_777023.1 | NCBI |
| *Callorhinchus milii* | Elephant Shark | ENaC $\alpha$ | JW872093.1* | NCBI |
| *Chrysemys picta bellii* | Western Painted Turtle | ENaC $\alpha$ | XP_005291372.1 | NCBI |
| *Gallus gallus* | Chicken | ENaC $\alpha$ | NP_990476.2 | NCBI |
| *Homo sapiens* | Human | ENaC $\alpha$ | NP_001029.1 | NCBI |
| *Hynobius nigrescens* | Sendai Salamander | ENaC $\alpha$ | BAI66492.2 | NCBI |
| *Latimeria chalumnae* | Coelacanth | ENaC $\alpha$ | H3AJ42 | UniProt |
| *Lethenteron camtschaticum* | Japanese Lamprey | ENaC $\alpha$ | APJL01005293.1* | NCBI |
| *Neoceratodus forsteri* | Australian Lungfish | ENaC $\alpha$ | H1AFJ5.1 | NCBI |
| *Petromyzon marinus* | Sea Lamprey | ENaC $\alpha$ | S4RTA3 | UniProt |
| *Protopterus annectens* | West African Lungfish | ENaC $\alpha$ | BAO27802.1 | NCBI |
| *Xenopus laevis* | African Clawed Frog | ENaC $\alpha$ | NP_001081392.1 | NCBI |
| *Erpetoichthys calabaricus* | Ropefish | ENaC $\alpha$ | XP_028666006.1 | NCBI |
| *Bos taurus* | Cow | ENaC $\beta$ | NP_001091544.1 | NCBI |
| *Callorhinchus milii* | Elephant Shark | ENaC $\beta$ | XP_007903981.1 | NCBI |
| *Chrysemys picta bellii* | Western Painted Turtle | ENaC $\beta$ | XP_005288958.2 | NCBI |
| *Gallus gallus* | Chicken | ENaC $\beta$ | XP_015149982.1 | NCBI |
| *Homo sapiens* | Human | ENaC $\beta$ | NP_000327.2 | NCBI |
| *Latimeria chalumnae* | Coelacanth | ENaC $\beta$ | H3AVV2 | UniProt |
| *Lethenteron camtschaticum* | Japanese Lamprey | ENaC $\beta$ | APJL01036227.1* | NCBI |
| *Neoceratodus forsteri* | Australian Lungfish | ENaC $\beta$ | H1AFJ6.1 | NCBI |
| *Petromyzon marinus* | Sea Lamprey | ENaC $\beta$ | S4RY81 | UniProt |
| *Protopterus annectens* | West African Lungfish | ENaC $\beta$ | BAO27803.1 | NCBI |
| *Xenopus laevis* | African Clawed Frog | ENaC $\beta$ | P51169 | NCBI |
| *Erpetoichthys calabaricus* | Ropefish | ENaC $\beta$ | XP_028670289.1 | NCBI |

**Supplementary Table 1** (continued)

| Species Name | Common Name | Protein | Accession Number | Database |
|---|---|---|---|---|
| *Bos taurus* | Cow | ENaC δ | XP_005217258.1 | NCBI |
| *Chrysemys picta bellii* | Western Painted Turtle | ENaC δ | XP_005293087.1 | NCBI |
| *Gallus gallus* | Chicken | ENaC δ | XP_004947475.1 | NCBI |
| *Homo sapiens* | Human | ENaC δ | AAI25075.1 | NCBI |
| *Latimeria chalumnae* | Coelacanth | ENaC δ | H3BHF6 | UniProt |
| *Xenopus laevis* | African Clawed Frog | ENaC δ | NP_001082645.1 | NCBI |
| *Bos taurus* | Cow | ENaC γ | NP_001180103.1 | NCBI |
| *Callorhinchus milii* | Elephant Shark | ENaC γ | XP_007903982.1 | NCBI |
| *Chrysemys picta bellii* | Western Painted Turtle | ENaC γ | XP_008163042.1 | NCBI |
| *Gallus gallus* | Chicken | ENaC γ | XP_015149986.1 | NCBI |
| *Homo sapiens* | Human | ENaC γ | NP_001030.2 | NCBI |
| *Latimeria chalumnae* | Coelacanth | ENaC γ | H3AU95 | UniProt |
| *Lethenteron camtschaticum* | Japanese Lamprey | ENaC γ | APJL01036229.1* | NCBI |
| *Neoceratodus forsteri* | Australian Lungfish | ENaC γ | H1AFJ7.1 | NCBI |
| *Petromyzon marinus* | Sea Lamprey | ENaC γ | S4RK61 | UniProt |
| *Protopterus annectens* | West African Lungfish | ENaC γ | BAO27804.1 | NCBI |
| *Xenopus laevis* | African Clawed Frog | ENaC γ | NP_001079123.1 | NCBI |
| *Erpetoichthys calabaricus* | Ropefish | ENaC γ | XP_028670288.1 | NCBI |
| *Lepisosteus oculatus* | Spotted Gar | γ-like | XP_006632023.1 | NCBI |
| *Scleropages formosus* | Asian Arowana | γ-like | XP_018588349.1 | NCBI |
| *Branchiostoma belcheri* | Lancelet | γ-like | XP_019619098.1 | NCBI |
| *Branchiostoma belcheri* | Lancelet | α-like | XP_019623744.1 | NCBI |

**Supplementary Table 2.** BayesTraits run parameters. [1]ML; Maximimum likelihood run. All ML runs included the commands: MLTries 10000; ScaleTrees 1000. [2]Markov chain Monte Carlo run. All MCMC runs included the commands: ScaleTrees 1000; Stones 100 10000. [3]Restricts reverse rate for trait 1 to 0. [4]Restricts trait 1 forward rates to be independent of trait 2, and trait 1 reverse rates to 0. [5]Restricts the equivalent site 1 and site 2 rates to be equal. [6]Restricts the equivalent site 1 and site 2 rates to be equal, but dependent on the status of the other site. [7]The AddTag and Fossil commands select and restrict a node given by the most recent common ancestor of the proteins specified. Nodes 1, 2, 3, and 4 in Fig. 2 are specified in commands as "ENaC", "bg", "alpha", and "ad", respectively.

| # | Trait 1 | Trait 2 | Model | Method | Commands |
|---|---|---|---|---|---|
| 1 | Terrestrial Lungs | Site 1<br>Site 2<br>PY<br>Lungs | Discrete:Independent | ML[1] | Res beta1 0[3] |
| 2 | Terrestrial Lungs | Site 1<br>Site 2<br>PY<br>Lungs | Discrete:Dependent | ML | Res q24 q13[4]<br>Res q31 q42 0 |
| 3 | Site 1 | Site 2 | Discrete:Independent | ML | Res alpha1 alpha2[5]<br>Res beta1 beta2 |
| 4 | Site 1 | Site 2 | Discrete:Dependent | ML | Res q12 q13[6]<br>Res q21 q31<br>Res q24 q34<br>Res q42 q43 |
| 5 | Site 1 | Site 2 | Discrete:Dependent *unrestricted ancestral nodes* | MCMC[2] | Commands from #4<br>Prior q34 exp 0.001<br>Prior q43 exp 0.001 |
| 6 | Site 1 | Site 2 | Discrete:Dependent *Trait 1 in ancestral nodes restricted to reflect divergent evolutionary model* | MCMC | Commands from #7<br>AddTag TENaC Cow_alpha Cow_beta<br>Fossil ENaC TENaC 15<br>AddTag Tbg Cow_beta Cow_gamma<br>Fossil bg Tbg 15<br>AddTag Talpha Cow_alpha SLamprey_alpha<br>Fossil alpha Talpha 15<br>AddTag Tad Cow_alpha Cow_delta<br>Fossil ad Tad 15 |
| 7 | Site 1 | Site 2 | Discrete:Dependent *Trait 2 in ancestral nodes restricted to reflect divergent evolutionary model* | MCMC | Commands from #7<br>AddTag TENaC Cow_alpha Cow_beta<br>Fossil ENaC TENaC 14<br>AddTag Tbg Cow_beta Cow_gamma<br>Fossil bg Tbg 14<br>AddTag Talpha Cow_alpha SLamprey_alpha<br>Fossil alpha Talpha 14<br>AddTag Tad Cow_alpha Cow_delta<br>Fossil ad Tad 14 |

| | | | | | |
|---|---|---|---|---|---|
| 8 | Site 1 | Site 2 | Discrete:Dependent<br>*Trait 1 in ancestral node 1 restricted to reflect convergent evolutionary model* | MCMC | commands from #7<br>AddTag TENaC Cow_alpha Cow_beta<br>Fossil ENaC TENaC 10<br>AddTag Tbg Cow_beta Cow_gamma<br>AddMRCA bg Tbg<br>AddTag Talpha Cow_alpha SLamprey_alpha<br>AddMRCA alpha Talpha<br>AddTag Tad Cow_alpha Cow_delta<br>AddMRCA ad Tad |
| 9 | Site 1 | Site 2 | Discrete:Dependent<br>*Trait 2 in ancestral node 1 restricted to reflect convergent evolutionary model* | MCMC | commands from #7<br>AddTag TENaC Cow_alpha Cow_beta<br>Fossil ENaC TENaC 11<br>AddTag Tbg Cow_beta Cow_gamma<br>AddMRCA bg Tbg<br>AddTag Talpha Cow_alpha SLamprey_alpha<br>AddMRCA alpha Talpha<br>AddTag Tad Cow_alpha Cow_delta<br>AddMRCA ad Tad |
| 10 | Terrestrial | PY | Discrete:Independent<br>*Unrestricted ancestral nodes* | MCMC | Commands from #1<br>PriorAll exp 0.001 |
| 11 | Terrestrial | PY | Discrete:Independent<br>*Trait 2 in ancestral nodes restricted to reflect divergent evolutionary model* | MCMC | Commands from #10<br>AddTag TENaC Cow_alpha Cow_beta<br>Fossil ENaC TENaC 14<br>AddTag Tbg Cow_beta Cow_gamma<br>Fossil bg Tbg 14<br>AddTag Talpha Cow_alpha SLamprey_alpha<br>Fossil alpha Talpha 14<br>AddTag Tad Cow_alpha Cow_delta<br>Fossil ad Tad 14 |
| 12 | Terrestrial | PY | Discrete:Independent<br>*Trait 2 in node 1 restricted to reflect convergent evolutionary model* | MCMC | Commands from #10<br>AddTag TENaC Cow_alpha Cow_beta<br>Fossil ENaC TENaC 11 |
| 13 | Terrestrial | PY | Discrete:Independent<br>*Trait 2 in node 2 restricted to reflect convergent evolutionary model* | MCMC | Commands from #10<br>AddTag Tbg Cow_beta Cow_gamma<br>Fossil bg Tbg 11 |

| | | | | | |
|---|---|---|---|---|---|
| 14 | Terrestrial | PY | Discrete:Independent<br>*Trait 2 in node 3 restricted to reflect convergent evolutionary model* | MCMC | Commands from #10<br>AddTag Talpha Cow_alpha SLamprey_alpha<br>Fossil alpha Talpha 11 |
| 15 | Terrestrial | PY | Discrete:Independent<br>*Trait 2 in node 4 restricted to reflect convergent evolutionary model* | MCMC | Commands from #10<br>AddTag Tad Cow_alpha Cow_delta<br>Fossil ad Tad 11 |

**Supplementary Table 3.** Primers for RT-PCR and expected product sizes.

| Target | | Primer sequence (5'–3') | Amplicon (bp) |
|---|---|---|---|
| *Erpetoichthys calabaricus* | | | |
| ENaC α | forward | ACATATTGCCTGGGGGCAC | 474 |
| | reverse | TTAGGCTCTTGGTGTCCATCTG | |
| ENaC β | forward | CCGGAATGGGTGTACTGCTA | 469 |
| | reverse | AAGGAGGAGGCCCCATGTAT | |
| ENaC γ | forward | AGCCGAACAACGAGCATGTA | 725 |
| | reverse | AGTTGCAAAGCTGGAAGCCT | |
| GAPDH | forward | TGAAAAGGCCTCTGCTCACC | 734 |
| | reverse | AGGTTTAACTGCGTCAGGGG | |
| *Xenopus laevis* | | | |
| ENaC α | forward | ACAGAGTGAGCCAGGATTGG | 628 |
| | reverse | AATTAACAGCTCCAGGTGGCA | |
| ENaC β | forward | CACATACCGCCGGCTCACT | 515 |
| | reverse | CGGCTTAACCCGTGAGTATTTGA | |
| ENaC γ | forward | GTCAGATCTCTGGGACAGATCCA | 847 |
| | reverse | GTTCCGAGCATCACAGGACA | |
| ENaC δ | forward | ACCACTTTCTGGCTTGTGCT | 320 |
| | reverse | CCTCCATTGACTTGGCCTGT | |
| β-actin | forward | GCCCGCATAGAAAGGAGACA | 610 |
| | reverse | GTCTGTCAGGTCACGTCCAG | |

**Tree File.**

```
#Nexus

Begin Trees;
     Translate
          1 Frog_alpha,
          2 Frog_beta,
          3 Frog_delta,
          4 Frog_gamma,
          5 Alungfish_alpha,
          6 Alungfish_beta,
          7 Alungfish_gamma,
          8 BCod_ASIC1,
          9 Arowana_g_like,
          10 Lancelet_a_like,
          11 Catfish_alpha,
          12 Chicken_alpha,
          13 Chicken_beta,
          14 Chicken_delta,
          15 Chicken_gamma,
          16 Coelacanth_ASIC1,
          17 Coelacanth_alpha,
          18 Coelacanth_beta,
          19 Coelacanth_delta,
          20 Coelacanth_gamma,
          21 Cow_alpha,
          22 Cow_beta,
          23 Cow_delta,
          24 Cow_gamma,
          25 Eshark_alpha,
          26 Eshark_ASIC1,
          27 Eshark_beta,
          28 Eshark_gamma,
          29 Elamprey_ASIC1,
          30 Human_alpha,
          31 Human_beta,
          32 Human_delta,
          33 Human_gamma,
          34 JLamprey_alpha,
          35 JLamprey_beta,
          36 JLamprey_gamma,
          37 JMedaka_ASIC1,
          38 Lancelet_g_like,
          39 Ropefish_beta,
          40 Ropefish_gamma,
          41 Ropefish_alpha,
          42 SLamprey_alpha,
          43 SLamprey_beta,
          44 SLamprey_gamma,
          45 Salamander_alpha,
          46 SpottedGar_g_like,
          47 WLungfish_alpha,
          48 WLungfish_beta,
          49 WLungfish_gamma,
```

```
            50 Turtle_alpha,
            51 Turtle_beta,
            52 Turtle_delta,
            53 Turtle_gamma
            ;
      Tree tree_1 =
((((10:0.240154,38:0.400628):0.26525,(9:0.332645,46:0.205856):0.672235):0.13
2642,(29:0.024507,
((8:0.020914,37:0.026452):0.020569,(26:0.041216,(11:0.054532,16:0.022955):0.
024701):1E-06):0.059287):
0.917353):0.065548,(((34:0.004582,42:0.050101):0.445435,((19:0.339745,(3:0.3
54015,((52:0.082886,14:
0.103927):0.085846,(32:0.199417,23:0.044511):0.204):0.090769):0.102711):0.15
4028,(25:0.80082,(41:
0.214639,((5:0.090541,47:0.111692):0.0775,(17:0.249053,(1:0.178596,(45:0.221
607,((50:0.067304,12:
0.056789):0.076495,(30:0.021723,21:0.082526):0.142622):0.057806):0.056807):0
.093321):0.04201):0.04742):
0.02314):0.039558):0.032187):0.182706,(((36:0.007161,44:0.030747):0.356102,(
(28:0.247241,40:0.284502):
0.040389,(20:0.138309,((7:0.113037,49:0.120798):0.151714,(4:0.176153,((53:0.
029715,15:0.054405):
0.057263,(33:0.014233,24:0.049296):0.128043):0.069185):0.046522):0.012495):0
.042939):0.250399):0.186618,((35:0.037746,43:1E-
06):0.193114,(39:0.259873,((18:0.135812,(27:0.296282,(6:0.093056,48:0.073352
):
0.216967):0.012272):0.028046,(2:0.20256,((51:0.013166,13:0.029852):0.064836,
(31:0.037238,22:0.057629):
0.087265):0.073248):0.067358):0.038976):0.116599):0.130724):0.134782):0.1477
99);
End;
```

**Supplementary Figure 1.** Multiple sequence alignment of proteins in Supplementary Table 1. Residues are colored by domain, as in Fig. 1: transmembrane and intracellular domains are blue, palm and β-ball domains are orange, finger domain is light green, GRIP domain is dark green, thumb domain is yellow-green, and the knuckle domain is brown. GRIP domain polybasic tracts and C-terminal PY motifs are underlined in red. Select conserved residues in ENaC subunits are bold.

```
Spotted Gar γ-like     ----------------------------------------MVISKKA------------------
Asian Arowana γ-like   ---------------------------------------------MPPKK---------------
Coelacanth ASIC1       -------------------MAAPCHSSDSV----------LSPYDSKDRE---------------
Catfish ASIC1          -------------------MVTLTTTREPV----------NNGSKAPKAG---------------
E. Lamprey ASIC1       ----------------------------------------MDLKGAPSDE---------------
E. Shark ASIC1         ------------------------------------------MDLKPPAE---------------
J. Medaka ASIC1        ----------------------MDLKADSD----------EMDYKRPAPI---------------
Black Rock Cod ASIC1   ----------------------MDLKADSE----------DMDYKRPAPI---------------
Lancelet γ-like        ------------------------------------------------MS---------------
Lancelet α-like        ----------------------------------------------MPRT---------------
E. Shark α             -----------------------------------------------------------------
Frog δ                 ---------------------------MES----------TEKEKKEGLI---------------
Coelacanth α           ---------------------------MSE----------KKEEKSKGLI---------------
S. Lamprey α           ---------------------PPTVLEFWR----------------GSVT---------------
J. Lamprey α           -----------------------------------------------------------------
W. Lungfish α          ----------------------MPNKEENA----------ENGKKKEGLF---------------
A. Lungfish α          ----------------------MTDKEEEA----------EGGKKKEPMI---------------
Ropefish α             -------------------------MSTTD----------DKQERKEGLL---------------
Frog α                 ---------------------------MTK----------EEKNEKEALI---------------
Salamander α           -----------------------MTEE---------------KKEKEGLI---------------
Cow α                  ---------------------MKGDKPEEPGPGPEPSGPPPPTEEEEALL---------------
Human α                -MEGNKLEEQDSSPPQSTPGLMKGNKREEQGLGPEPAAPQQPTAEEEALI---------------
Chicken α              -MGTASRGGSVKAEKMPEGEKTRQCKQETE--------QQQKEDEREGLI---------------
Turtle α               ----MHQVVAVKAENVPVGERLRRCKQEAEKQQKVEEVAEKLEKECQGLI---------------
Cow δ                  --------MENGRLMAQVGRPGWGQAKWWAGAPLSLTSQMQAEGTGQTVGGGPGTWTCPQASPPT
Human δ                MAEHRSMDGRMEAATRGGSHLQAAAQTPPRPGPPSAPPPPPKEGHQEGLV---------------
Coelacanth δ           -----------------------CLQSLKM---------AQEEDKEEAVI---------------
Chicken δ              -----------------------MEQEAAR----------EEEERKEGLI---------------
Turtle δ               ------------------------MEQEWV----------NEEMGDEGLI---------------
S. Lamprey β           ----------------------MKIRKYLT----------RSLHRLQ-KG---------------
J. Lamprey β           -------------------MANMKIRKYLT----------RSLHRLQ-KG---------------
E. Shark β             -------------------MLWVSARRCLS----------QALRRLQ-DG---------------
Coelacanth β           ----------------------MSVRKYFT----------RALHRLQ-KG---------------
Ropefish β             ----------------------MGVQKYLT----------YALHRIQ-KG---------------
W. Lungfish β          ---------------------MSFLKRFCV----------RSWHRIK-KG---------------
A. Lungfish β          ----------------------MFLKRWFI----------RALHRLQ-KG---------------
Frog β                 ----------------MIHGKMKRLKRYFT----------RALHRIQ-KG---------------
Cow β                  ----------------------MHVKKYLL----------KGLHRLQ-KG---------------
Human β                ----------------------MHVKKYLL----------KGLHRLQ-KG---------------
Chicken β              ----------------------MNLKRYFV----------RALHRLQ-KG---------------
Turtle β               ----------------MFTGMTMNFKRYFI----------RVLHRLQ-KG---------------
S. Lamprey γ           ------------------MASEGDSKRVLH----------RVKDTLKIDG---------------
J. Lamprey γ           -----------------MASEGDSNKRVLH----------RVKDTLKIEG---------------
Ropefish γ             --------------------MESVAKKLPK----------KVKEKLPVTG---------------
E. Shark γ             --------------MESVGAMEPGKRKLSA----------KIMEKLPVTG---------------
W. Lungfish γ          ---------------------MKNTKKFKE----------SVKKQLPLTG---------------
A. Lungfish γ          ---------------------MGHGRRISE----------SIKKQLPVTG---------------
Coelacanth γ           ------------------ATMTSRKKSLPE----------KIKENLPVTG---------------
Frog γ                 --------------------MSKSGKKLTQ----------KLKKNLPVTG---------------
Cow γ                  ---------------------MAPGEKIKA----------KIKKNLPVTG---------------
Human γ                ---------------------MAPGEKIKA----------KIKKNLPVTG---------------
Chicken γ              ----------------------MAPGKITA----------RIKKTLPVRG---------------
Turtle γ               --------MEPPRPGDTRELNMAPGKTIKA----------KIKKTLPVTG---------------
```

| | |
|---|---|
| Spotted Gar γ-like | ---------------TLQSICRETLIHTSA**HG**VSSILRS-RSNHQKNCWIIFVVVVVGCML-**W**QC |
| Asian Arowana γ-like | -----------PGAMTSLWMLRDDIQHTTA**HG**IPNIFRA-RHWFRRSLWAMFVIFAFCCAI-**W**QC |
| Coelacanth ASIC1 | -----------KRNHSLKQITIAFVKNSKF**HG**IRYIFAYHISKQRRAIWFLAFFIATGLLAI**W**SL |
| Catfish ASIC1 | ----------FERMSSMAKITLAFVFRTKV**HG**LRYVFAADKSKPRRFFWLVAICVCLALLFI**W**SC |
| E. Lamprey ASIC1 | -----------SLDQARPSSVATFADSCTL**HG**IRHIFSPGGLSVRRLLWLLAFLGSLSLLV-**L**QS |
| E. Shark ASIC1 | -----------DGVSNHPASVEAFAKTSTL**HG**ISHIFTYERFSFKRIIWTLAFLGSLSFLV-**H**TC |
| J. Medaka ASIC1 | ---------------------EVFATRSTL**HG**ISHMFTYERMCLKRTLWILFFMLSVGVLV-**M**VC |
| Black Rock Cod ASIC1 | ---------------------EVFASRSTL**HG**ISHMFTYERMCIKRTLWILFFLSSVGVLV-**M**VC |
| Lancelet γ-like | -----------EKRPSVRSTLRKYGENTSA**HG**IPRAVTT-KSLPRRLFWTCLFLASFSYFL-**Y**QA |
| Lancelet α-like | -----------TDNKVASALLE-FADTTTT**HG**VPRAVGS-SSLLRKICWTVAFVASLGYFL-**Y**QA |
| E. Shark α | --------------MDLARALALSRREAIC**HP**APRKP-------------------------**W**R- |
| Frog δ | -----------EFYDSFEDMLTFFCDNTTI**HG**TVRLNCSRKNKMKTTFWLVLYFVSFAMMY-**W**QF |
| Coelacanth α | -----------EFYSSYSDLFQFFCSTTTI**HG**AIRLVCTERNKMKTAFWSMLFVASFGLMY-**W**QF |
| S. Lamprey α | -----------DFYDSYDEMFEFFCDNTTI**HG**TIRLVCSKRNKLKTAFWSLLFTVTVILFY-**Y**TS |
| J. Lamprey α | -------------------MFEFFCDNTTI**HG**AIRLVCSKRNKLKTAFWSLLFIVTVILFY-**Y**TS |
| W. Lungfish α | -----------EFYDSFQELFEFFCINTTI**HG**TIRMVCSKHNNMKTAFWTILFIATFGIMY-**W**QF |
| A. Lungfish α | -----------GFYDSYQELFEFFCNNTTI**HG**TIRMVCSKHNNMKTVSWTILFITTFGVMY-**W**QF |
| Ropefish α | -----------EFYTSYSDLFNFFCSNTTI**HG**AIRLVCSENNRMKTAFWAILFPGTVAILY-**W**QF |
| Frog α | -----------EFFSSYRELFEFFCSNTTI**HG**AIRLVCSRRNRMKTAFWLVLFLVTFGLMY-**W**QF |
| Salamander α | -----------EFYSSYRELFEFFCNNTTI**HG**AIRLVCSAHNRMKTAFWVVLFIASFGLLY-**W**QF |
| Cow α | -----------EFHRSYRELFEFFCNNTTI**HG**AIRLVCSQHNRMKTVFWAVLWLCTFGMMY-**W**QF |
| Human α | -----------EFHRSYRELFEFFCNNTTI**HG**AIRLVCSQHNRMKTAFWAVLWLCTFGMMY-**W**QF |
| Chicken α | -----------EFYGSYQDVFQFFCSNTTI**HG**AIRLVCSKKNKMKTAFWSVLFILTFGLMY-**W**QF |
| Turtle α | -----------EFHKSYHELFQFFCNNTTI**HG**AIRLVCSKRNKMKTAFWSVLFFLTFGLMY-**W**QF |
| Cow δ | LPEEEHGERLVELHASFRELVTFFCTNSTI**HG**TIRLVCSSQNRLKTASWGLLLAGALGVLY-**W**QF |
| Human δ | -----------ELPASFRELLTFFCTNATI**HG**AIRLVCSRGNRLKTTSWGLLSLGALVALC-**W**QL |
| Coelacanth δ | -----------EFYDSFKDLFQFFCAHTTV**HG**GIRLICSERNMMKTAFWIILFFASFGLMY-**W**QF |
| Chicken δ | -----------EFYDSFKDMFEFFCKNTTI**HG**TIRLVCSSSNKMKTAFWTLLLLASFGMLY-**W**QF |
| Turtle δ | -----------EFYSSFKDMFEFFCKNTTI**HG**TVRLVCSSSNRMKTAFWTLLFLASFGMLY-**W**QF |
| S. Lamprey β | -----------PVA-SVSELLYWYCMNTNT**HG**CKRIVVY--GKKKRVLWFLITIIMLGVVL-**W**QW |
| J. Lamprey β | -----------PVA-SVSELLYWYCMNTNT**HG**CKRVVVY--GKKKRVLWFLITIIMLGVVL-**W**QW |
| E. Shark β | -----------PGE-SYRELLVWYCETTST**HG**PKRILTE--GPKKRALWLLLTLLLGGVVC-**W**QW |
| Coelacanth β | -----------PGY-TYKELLVWYCDNTNT**HG**PKRIIKE--GPKKQVLWFILTLTFTALIF-**W**QW |
| Ropefish β | -----------PGY-TYKELLVWYCNNTNT**HG**PKRIVTE--GPKKRFLWFLFTLVFAALVF-**W**QW |
| W. Lungfish β | -----------PGY-GYAELFHWYCDNTNT**HG**PKRLIIE--GPKKKAMWGLLTITFACLVF-**W**NW |
| A. Lungfish β | -----------PGY-GYSELFVWYCNNTNT**HG**PKRLIIE--GPKKKTLWSLFTVTFACLVF-**W**QW |
| Frog β | -----------PGY-TYKELLVWFCDNTNT**HG**PKRIIKE--GPKKRVMWFILTLVFAGLVF-**W**QW |
| Cow β | -----------PGY-TYKELLVWYCDNTNT**HG**PKRIICE--GPKKKAMWFVLTLLFTSLVC-**W**QW |
| Human β | -----------PGY-TYKELLVWYCDNTNT**HG**PKRIICE--GPKKKAMWFLLTLLFAALVC-**W**QW |
| Chicken β | -----------PGY-TYKELLVWYCDNTNT**HG**PKRIIKE--GPKKKVMWFFLTLLFASLVF-**W**QW |
| Turtle β | -----------PGY-TYKELLVWYCDNTNT**HG**PKRIIRE--GPKKKVIWFFLTLLFASLVF-**W**QW |
| S. Lamprey γ | -----------PDP-SITDLLDFYLNNTNM**HG**MRRIAVS-KGPIKKTIWIVFSLIAVAMVF-**W**QG |
| J. Lamprey γ | -----------PDP-SITDLLDFYLNNTNM**HG**MRRIAVS-KGPIKKTIWIVFSLIAVAMVF-**W**QG |
| Ropefish γ | -----------PYAITVKELMVWYCNYTNT**HG**CRRIVVS-RGRLRRWIWTVLTLSAVALIS-**W**QC |
| E. Shark γ | -----------PQALSMSELARWYCYNTNT**HG**FRRIVVS-RGRLRRGAWVLLTGCAASLIV-**W**QC |
| W. Lungfish γ | -----------PESRTVKDLMDWYCNNTNT**HG**CRRIAVS-RGHLRRWIWICFTLTAVAIIF-**W**QY |
| A. Lungfish γ | -----------PEAPTVKNLMDWYLNNTNT**HG**CRRIAVS-RGYLRRWIWICFTVSSVGMIF-**W**QW |
| Coelacanth γ | -----------PQALSISELMRWYCYNTNT**HG**CLRIVAS-RGRLRRWIWILLTLSAVALIF-**W**QC |
| Frog γ | -----------PQAPTLYELMQWYCLNTNT**HG**CRRIVVS-KGRLRRWIWISLTLCAVAVIF-**W**QC |
| Cow γ | -----------PQAPNIKELMQWYCLNTNT**HG**CRRIVVS-RGRLRRLLWILFTLTAVALIF-**W**QC |
| Human γ | -----------PQAPTIKELMRWYCLNTNT**HG**CRRIVVS-RGRLRRLLWIGFTLTAVALIL-**W**QC |
| Chicken γ | -----------PQAPTLRELMRWYCLNTNT**HG**CRRIVVS-RGRLRRFIWILLTLSAVGLIL-**W**QC |
| Turtle γ | -----------PQAPTVSELMHWYCMNTNT**HG**CRRIVVS-RGRLRRFIWILLTLSAVGLIL-**W**QC |

| | |
|---|---|
| Spotted Gar γ-like | SELINTFFHYPSQEKVTLVNSARLK-**FPAVT**F**CN**L**N**QVRKSLMLSKFSFLK-------GGLYFLN |
| Asian Arowana γ-like | MEIIMTFYSYPSHEKIRLISDTKLM-**FPAVT**I**CN**L**N**SVRHSALKRNFALNNTFLDFCL------- |
| Coelacanth ASIC1 | NRILYLFS-YPAVIKMQMIWARHLY-**YPTVT**I**CN**Y**N**LFRLSRMTK-------------ADLYYSG |
| Catfish ASIC1 | NRLL-YLLSFPAVTKIYMVWANNMT-**FPAVT**L**CN**Q**N**LFRVSSLTK-------------ADLYHSG |
| E. Lamprey ASIC1 | LDWVQYYLRYPCVTKQDEVSTPLTV-**FPAVT**L**CN**L**N**EFRFSRMTR-------------NDLYHAG |
| E. Shark ASIC1 | TQRIQYYFQYPHVTKLDEISAANMT-**FPAIT**I**CN**L**N**EFRFSKITK-------------NDLYHAG |
| J. Medaka ASIC1 | VDRVQFYFEYPHVTKLDEVAASMIV-**FPAIT**F**CN**L**N**SFRFSRVTR-------------NDLYHAG |
| Black Rock Cod ASIC1 | VDRVQLYFQYPHVTKLDEVSAPMMV-**FPSVT**F**CN**L**N**SFRFSRVTR-------------NDLYHAG |
| Lancelet γ-like | QTLVNKYLVYPVNTDVK-IEWSELE-**FPAVT**I**CN**A**N**PLRYRELIKR----------------GSA |
| Lancelet α-like | SMLFNKYFDYPVATDIS-IKFATIE-**FPAVT**I**CN**L**N**PVRLSKLNTAGGEFSSY---IISDVGTTA |
| E. Shark α | ---------FPVKSHRCPPSFPLLLS**LPPSP**S**PS**P**S**PFTHSP----------------------- |
| Frog δ | GQLTDQYWAYPTSTIIG-LQSKGKI-**FPAVT**I**CN**L**N**PYRFDQVNMYINQLDQLANETLYSLYEYR |
| Coelacanth α | GIIFGHYFSYPVSMSLT-LEHKKLL-**FPAVT**V**CT**L**N**PYRYKEVESELKELDSLTADTLFELYRYN |
| S. Lamprey α | ALVFLQYYSYTVAVTMG-LMFQQST-**FPAIT**V**CS**L**N**PYRYEVVQSSLSQLDSMTGQALQQLYGYQ |
| J. Lamprey α | ALVFLQYYSYTVAVTMG-LMFQQST-**FPAIT**V**CS**L**N**PYRYEAVQSSLRELDSMTGQALHRLYGYQ |
| W. Lungfish α | GLLLDQYYSFPVSITMA-VNYDKLV-**FPAVT**V**CT**L**N**PYRYNAVSTELANLDCYTEQLLSTLYHYT |
| A. Lungfish α | GLLLGQYYSYPVSITMS-VNFDKLI-**FPAVT**V**CT**L**N**PYRYNVVSTQLANLDCYTEELLSTLYHYN |
| Ropefish α | GLLFGQYFSHPVSIGVS-VNFNELQ-**FPSVT**V**CT**L**N**PYRYSAVREELKELDAVTEETLYKLYGYT |
| Frog α | GLLFGQYFSYPVSINLN-VNSDKLP-**FPAVT**V**CT**L**N**PYRYKAIQNDLQELDKETQRTLYELYKYN |
| Salamander α | GLLFGQYFSYPVSINMN-VNSDKLL-**FPAVT**V**CT**L**N**PYRYTAVLEDLRELDRLTEQTLYDLYRYN |
| Cow α | GQLFGEYFSYPVSLNIN-LNSDKLV-**FPAVS**I**CT**L**N**PYRYKEIQEELEELDRITEQTLFDLYKYN |
| Human α | GLLFGEYFSYPVSLNIN-LNSDKLV-**FPAVT**I**CT**L**N**PYRYPEIKEELEELDRITEQTLFDLYKYS |
| Chicken α | GILYREYFSYPVNLNLN-LNSDRLT-**FPAVT**L**CT**L**N**PYRYSAIRKKLDELDQITHQTLLDLYDYN |
| Turtle α | GILYRQYFSFPVNLNLN-LNSDRLT-**FPAVT**L**CT**L**N**PYRYSAVQKELDELDRITHQTLMDLYNYN |
| Cow δ | ALLFEQYWRYPVIMTVS-VHSERKL-**FPSVT**L**CD**M**N**PHRPHLARHHLRVLDDFARESIYSLYRFN |
| Human δ | GLLFERHWHRPVLMAVS-VHSERKL-**LPLVT**L**CD**G**N**PRRPSPVLRHLELLDEFARENIDSLYNVN |
| Coelacanth δ | GLLFSQYWGYPVSVAIR-VHSGPKI-**FPAVT**V**CT**L**N**PYRYTQVHKYLKELDQMALEVLSTWYGFN |
| Chicken δ | ALMFSQYWDYPVVLTMS-MHSEPKM-**FPAIT**I**CN**L**D**PYRFDLVSEHLAQLDRMAEKSVTVLYGIN |
| Turtle δ | ALLFSQYWTYPVIMTMS-VHSEPKM-**FPAIT**L**CN**L**D**PYRFDLVSERLAQLDQMAEEAIANLYGYK |
| S. Lamprey β | VLLFQAYLSYGVSVSVN-MGFQRMN-**FPAVT**V**CN**L**N**AYRYSSMKDKIKDLEAYTRVALQTLYNYT |
| J. Lamprey β | VLLFQAYLSYGVSVSVN-MGFQRMN-**FPAVT**V**CN**L**N**AYRYSSIKDKIKDLEAYTRVALQTLYNYN |
| E. Shark β | GVLVQRYLSGETISTLR-TGFKAMV-**FPAVT**L**CN**V**N**PFRYSRSRGLLQPLDRLAELALQRIYMYN |
| Coelacanth β | GLLIQTYLSYGVSTSLS-MGFRAME-**FPAVT**V**CN**V**N**PMKYSEPPKLIQSMHFLIFLFLTKKISKR |
| Ropefish β | GILIQTYMSWGVSTSLS-VGFKTAP-**FPAVT**I**CN**V**N**PFKYSKVKPLIEDLDEAARTALAKIHTYF |
| W. Lungfish β | GVLIQTYLSWGVSVSLS-VGFSSLA-**FPAVT**I**CN**S**N**PFKYSRIKPLLTELDGFAASLLERIYIYS |
| A. Lungfish β | GLLIQTYLSWGVSVSLS-VGFRGMD-**FPAVT**V**CN**V**N**PFKYSKVKPLLKELDELVDILLEQFYSYS |
| Frog β | GVLILTYLSYGVSVSLS-IGFKTME-**FPAVT**L**CN**A**N**PFKYSRVKPLLKELDELVATALDRIQFSS |
| Cow β | GLFIKTYLNWEVSVSLS-IGFKTMD-**FPAVT**I**CN**A**S**PFQYSKVQHLLKDLDELMEAVLGRILGPE |
| Human β | GIFIRTYLSWEVSVSLS-VGFKTMD-**FPAVT**I**CN**A**S**PFKYSKIKHLLKDLDELMEAVLERILAPE |
| Chicken β | GILINTYLSYNVTSSLS-IGFKTMK-**FPAVT**V**CN**A**N**PFKYSEVRPLLKELDKLIEAALERILQPT |
| Turtle β | GILIDTYLSYSVSSSLS-IGFKTMK-**FPAVT**V**CN**A**S**PYKYSKVRHLLKELDELTEAALERILQSK |
| S. Lamprey γ | IQLIQSF--YSIAVSVT-INYQKLP-**FPAIT**V**CS**L**N**PYKYNQSQALLEKLDRNTAVALHNIGIAV |
| J. Lamprey γ | IQLIQSF--YSIAVSVT-INYQKMP-**FPAIT**I**CS**L**N**PYKYNQSQALLEKLDRNAAVALHNIGIAV |
| Ropefish γ | ALLIQTY--YSSSVSVT-VQFQTLT-**FPAVT**V**CN**L**N**PLRYSATKQLLTELDEQAERALQELYSPK |
| E. Shark γ | ALLASAY--YTVSVSIT-VHFQELP-**FPAIT**I**CN**I**N**PYRYSATRWLVGELEKATLTVLDELYKYT |
| W. Lungfish γ | TLLVMSY--YSVTVSVM-VKYQTLS-**FPAVT**V**CN**I**N**PYKPNTTISLLDELNRQARKILEKLYGFC |
| A. Lungfish γ | TLLLMSY--YTVSVSVT-VQFQTLP-**FPAVT**I**CN**I**N**PYRKNATSALLEELDKQTKLILKELYTSC |
| Coelacanth γ | ALLIISY--YSVTVAVS-VQFQELN-**FPAIT**I**CN**I**N**PYRYSATGELLQELERETKNALKVLYDFP |
| Frog γ | ALLLMSY--YSVSASIT-VTFQKLV-**YPAVT**I**CN**L**N**PYSYSKVKDRLAALEKETSQTLKNIYGFT |
| Cow γ | ALLISSF--YTVSVSIK-VHFQKLD-**FPAVT**I**CN**I**N**PYKYSAVRHLLADLEQETRAALKTLYGFS |
| Human γ | ALLVFSF--YTVSVSIK-VHFRKLD-**FPAVT**I**CN**I**N**PYKYSTVRHLLADLEQETREALKSLYGFP |
| Chicken γ | AELLLNY--YSASVSVT-VQFQKLP-**FPAVT**I**CN**I**N**PYKYSSMKDYLSELDKETKKALETFYGFS |
| Turtle γ | AELIMSY--YTASVSVT-VQFQKLP-**FPAVT**I**CN**I**N**PYKYSAMKEHLSELDKETKNALETLYGFS |

| | |
|---|---|
| Spotted Gar γ-like | ----------------------------------------------------------------- |
| Asian Arowana γ-like | ----------------------------------------------------------------- |
| Coelacanth ASIC1 | -----------------------------YWLDLLHQDLSVNQDSLGVL---------------- |
| Catfish ASIC1 | -----------------------------YWIDIMHANHSVNRQSMAMLK--------------- |
| E. Lamprey ASIC1 | -----------------------------ELLALLDERMEIVEPRFADAQVIAQLRK-------- |
| E. Shark ASIC1 | -----------------------------ELLTLLNNRYEIPDPHLAERHILEALVE-------- |
| J. Medaka ASIC1 | -----------------------------ELLALLNGRYEIRDPHLVEENVLQVLRE-------- |
| Black Rock Cod ASIC1 | -----------------------------ELLALLNGRYEIRDPHMVEEHVLQILKE-------- |
| Lancelet γ-like | AFQNAGFIPKPQP----------------QGQNNPNSNGSTPATGSTTVATAPTTTQNSTS---- |
| Lancelet α-like | PPSPGPRRKRGVDTSQDNRPEYDV-----DWEDPYPPREESAEEHELLRERRETGSSAGSADSQG |
| E. Shark α | ----------------------------------------------------------------- |
| Frog δ | APESGQQVVDLQD-LLNNLTGQVN-GGFYLDESIVLLKLQENGSGPALPG--------------- |
| Coelacanth α | TSQHGTSDDSMQSDIRRSNRIILS-----APDRVPLQVLDEPAAEHARTENQMAGT--------- |
| S. Lamprey α | PPATKAAGTAAAP----------------GIRLDTGVVLERTGPDT------------------- |
| J. Lamprey α | PPATQSAGTAAAP----------------GIRLDTGVVLERTGPDT------------------- |
| W. Lungfish α | PANSNQSACNNTS-NKQDS----------NNKYIPLEFLTYNDTQSGYPFKGTTNGSSSFN---- |
| A. Lungfish α | PLTSGNQSACNSS-STAGTRAFDE-------SYMKLEFLNDENTAYSGPVKGATNSTSPVN---- |
| Ropefish α | FSKKVQQNSTSHT-SATEKSRNSN-PL--FNKKFVLEVLNRETSVNFNTPEKKGNHGV------- |
| Frog α | --------STGVQ-GWIPNNQRVKRDR--AGLPYLLELLPPGSETHRVSRSVIEEELQVK----- |
| Salamander α | SSGMDALYHESRRER--------------RSATFPLFPLERLSPEQGAGGRVRRSSGAAVREEEL |
| Cow α | -------SSKTLV-AHARSRRDLR-----EPLPHPLQRLPVPAPPHAARGVRRAGSSM------- |
| Human α | -------SFTTLV-AGSRSRRDLR-----GTLPHPLQRLRVPPPPHGARRARSVASSL------- |
| Chicken α | MSLARSDGSAQFS-HRRTSRSLLH-----HVQRHPLRRQKRDNLVSLPENSPSVD---------- |
| Turtle α | MSLVQSNWAAQSS-RKRSPRSLSH-----HVHRHPLRRHKRDEPASLKGNSPPVD---------- |
| Cow δ | FSDSMDALGAEVP----GPEPAFH-----LDRRIRLQRLRPLDGQNR------------------ |
| Human δ | LSKGRAALSATVP----RHEPPFH-----LDREIRLQRLSHSGSRVR------------------ |
| Coelacanth δ | ASEDITPNDSIGDGAGHGIKNISDNMNITLDQSIPLVLIRDKDSLDSSSAHSFPLD--------- |
| Chicken δ | TSASLFHVNEKSI-HVRDLPSTGNHNGSSFKLSQKFSLLRTTEFNNRT----------------- |
| Turtle δ | TSIFSSRYKKDIS-VKGGSANLSS-SSFQLNRHISLVMLKEPDTGSR------------------ |
| S. Lamprey β | DSSTPSAYDTSYAVG--------------PWQEIPLVLIDRRDPNRTVVTEVMCSRAAVGIETHT |
| J. Lamprey β | DSSTPSPYNTSYA-KA-------------PWQDIPLVLIDRRDPNRTVVTEVVSKQSYLINGSID |
| E. Shark β | QNRTLPPPMDDLEDKWSRGGAGLG-----PGPLVPLVIIEQTEKGEEVLVRILGDAESDVP---- |
| Coelacanth β | ERHSLTPVWKTNL-KKLLILPKTSQACVYLQKPQFLYLVSFFNKYHILKIVVLPL---------- |
| Ropefish β | ISGKVPELPLMA-----NTSLVSV-----FLQNIPLVIMDESDVNNPAIISLFENGTDGF----- |
| W. Lungfish β | QSGTLPDPLLDTT-SRNATGQNL------LWYHLPLVIIDETDGDNPVIINILGPNNLTSTNGTT |
| A. Lungfish β | TNGTLPVVFPDMR-SS-YLTGDPP-----PWYQIPLVMIDETDADNPTVTNVLGTDAL------S |
| Frog β | QNQGNTFTHNNQT-RQ-NVTLDPA-----LWNHIPLVVIDETDPRNPIIHNIFDNNAVYSKN--S |
| Cow β | LSQVNDTRALNLS----------------IWHHTPLVFINEQNPHHPVVLDLFEDNFNGSASNSP |
| Human β | LSHANATRNLNFS----------------IWNHTPLVLIDERNPHHPMVLDLFGDNHNGLTSSSA |
| Chicken β | HGDPISPLLLNNS-NA-TEGLDLD-----LWNQIPLVLIDEQDKDNPVIVEIFETNQSAAGNQ-T |
| Turtle β | HGATISAPPLNSS-ETVSQQLDLK-----LWNQIPLVLIDESDPDHPVIIDLFETDQSGSGTRPN |
| S. Lamprey γ | TNLSAAKRDD-------------------EPLPIPLVWLDTTVTNQTVVTDVISGKFHVVPGKVE |
| J. Lamprey γ | TNLSAAKRDD-------------------EPLPMPLVWLDTTVTNQTVVTDVISGKFHVVPGEVE |
| Ropefish γ | EPTEKRKRRGLVTDKHAEDGQVVSN----LLKDIPLFWLDKKTFSAAFAPN---NRKRTYPEKFL |
| E. Shark γ | DAEDDGGEVRPQH-E-AEPGEGSR-PS--LFQNIPLLQIQARGPEYSIVSNLLSRRQHRVNGTVS |
| W. Lungfish γ | TDCGSGNFSSRSL-SLDDIPVKTRSLN--LLQDMSLIKVETAKDGQLVASDIVTDLQYRISGTAI |
| A. Lungfish γ | TGCSNRKLRSVLL-NEAPEEDSGV-AK--LLQDMPLMKFEVIKEDHVIVSELSSNRQYRINNTFI |
| Coelacanth γ | LDDNESHVLRSTE-TSLNSESDKE-VL--FSRSLPLLKIEEMEQNYTIVSDVFSDVKQRVNAPLM |
| Frog γ | EPLIRSKRDVGVNVENSTEDI--------FLKQIPLYRLESVKGSQLVVSDLKTKKRTRMSAKVI |
| Cow γ | EITSRKRREAQSW-SSVRKGTDPK-----FLNLAPLMAFEKGDTGKA--RDFFTGRKRKVNARII |
| Human γ | E--SRKRREAESW-NSVSEGKQPR-----FSHRIPLLIFDQDEKGKA--RDFFTGRKRKVGGSII |
| Chicken γ | EGKTKVRRAAGDWNGTESL----------FFRHVPLLRFE---NSFRAATDLRSGRKRKVEGSVF |
| Turtle γ | EGKSKVRRDADDWNATESK----------FFEIIPLLKFE--DLSKKTVTEIPSGNKRKIETSVF |

| | |
|---|---|
| Spotted Gar γ-like | ------------------------------------TDYANS**S**Q**N**QKAY---------------- |
| Asian Arowana γ-like | ---------------------AQRWNSTGYPG**A**---TRDTN-R**C**A-NSF--------------S- |
| Coelacanth ASIC1 | ---------------------REDTKQKILQL**S**-----------NFTQY--------------TP |
| Catfish ASIC1 | ---------------------HSRHRERLMRL**L**-----------DFSDY--------------VP |
| E. Lamprey ASIC1 | ------------------------------------------H**A**DFRVH--------------KP |
| E. Shark ASIC1 | ------------------------------------------K**A**NFRNF--------------KP |
| J. Medaka ASIC1 | ------------------------------------------R**A**DFESY--------------KP |
| Black Rock Cod ASIC1 | ------------------------------------------R**A**NFDNY--------------KP |
| Lancelet γ-like | ------------------------------------DEDDD-D**Y**DYDGY--------------HG |
| Lancelet α-like | SSDSSDSSDSSD---------SSDSSDSSDSS**D**SSDSSDSSDS**S**DYASYMYGGTWSSHYGQDGSS |
| E. Shark α | ---------------------------------------------------------------SP |
| Frog δ | ---------------------EKKFK-VGFKL**C**---NSSRD-D**C**YYKVF--------------WS |
| Coelacanth α | ----DINNPALY---------KGEFRKIGFKL**C**---NASGL-S**C**FYQAY--------------SS |
| S. Lamprey α | ---------------------------VGFKL**C**---NATGG-D**C**FYQSY--------------GS |
| J. Lamprey α | ---------------------------VGFKL**C**---NATGG-D**C**FYQSY--------------GS |
| W. Lungfish α | ---------------------NSEFYRVGFKV**C**---NDTDG-V**C**FYQIY--------------SS |
| A. Lungfish α | ---------------------HTEFYRIGFKL**C**---NATGE-D**C**FYQTY--------------SS |
| Ropefish α | ----AKNNPPLH---------NQNWQ-IGFKL**C**---NASGQ-D**C**YFQAY--------------SS |
| Frog α | ---------------------RREWN-IGFKL**C**---NETGG-D**C**FYQTY--------------TS |
| Salamander α | PID------------------SISWN-IGFKL**C**---NDSGK-D**C**FYQKY--------------SS |
| Cow α | ----RDNNPQVN---------RKDWK-IGFQL**C**---NQNKS-D**C**FYQTY--------------SS |
| Human α | ----RDNNPQVD---------WKDWK-IGFQL**C**---NQNKS-D**C**FYQTY--------------SS |
| Chicken α | ---------------------KNDWK-IGFVL**C**---SENNE-D**C**FHQTY--------------SS |
| Turtle α | ---------------------KSDWK-IGFIL**C**---NETNE-D**C**FHQTY--------------SS |
| Cow δ | ---------------------------VGFKL**C**---NSTGG-D**C**VERAY--------------SS |
| Human δ | ---------------------------VGFRL**C**---NSTGG-D**C**FYRGY--------------TS |
| Coelacanth δ | ---------------------KKGFR-VGFRL**C**---NVTGK-D**C**FYQSY--------------SS |
| Chicken δ | ---------------------GKRQSLVGFRL**C**---NATGG-N**C**FYKTY--------------SS |
| Turtle δ | ---------------------KKHFK-VGFKL**C**---NATGG-N**C**FYKAY--------------SS |
| S. Lamprey β | VDNRVF---------------HIGFCKSALGV**C**---CDSAGDK**C**FYSEY--------------LS |
| J. Lamprey β | TVSVPPERAAADATSSLIPHLGKDVR-IGFRL**C**---DTAGD-K**C**FYSEY--------------LS |
| E. Shark β | ----RDAKLGRG---------PRSYK-VALHL**C**---SKDGR-D**C**LYRNF--------------TT |
| Coelacanth β | ---------------------ANEKKQKVSVL**C**KDERMGKKKK**S**SVYFT--------------NA |
| Ropefish β | ----VQNNPHPP---------TSDLK-VAIKL**C**---NANKT-E**C**LYWNF--------------TS |
| W. Lungfish β | AGYPIP---------------PRRYK-VAFEL**C**---NASGS-D**C**FYKNF--------------SS |
| A. Lungfish β | PTNNSTTNSSTE---------ARRYK-VAFHL**C**---NTNGT-D**C**FYKNF--------------SS |
| Frog β | SIRNSSEDQTSY---------SQRYK-VAMKL**C**---TNNNT-Q**C**VYRNF--------------TS |
| Cow β | APGRPC---------------SAHRCKVAMRL**C**---SHNGT-T**C**TFRNF--------------SS |
| Human β | SEKIC----------------NAHGCKMAMRL**C**---SLNRT-Q**C**TFRNF--------------TS |
| Chicken β | AAPPAPANVTSE---------EKKYK-LAVKL**C**---SHQGSNN**C**TYRNF--------------TS |
| Turtle β | NSSPALSNVTSE---------VKKHK-VAVKL**C**---HHKDIPQ**C**MYWNF--------------TS |
| S. Lamprey γ | MRSYFSQNY------------QSSEPLIAIEV**C**---GEER--K**C**IYNAF--------------TS |
| J. Lamprey γ | MRSYFSQNY------------QSSEPLIAIEV**C**---GEEK--K**C**IYNAF--------------TS |
| Ropefish γ | SKSGSRLRF------------HEAQRQAGFQL**C**---NSTNVTD**C**VVYAF--------------DT |
| E. Shark γ | THSLGNADILNQ---------EHL---VGFKL**C**EGGDIDSS-D**C**TIYTF--------------TS |
| W. Lungfish γ | TRMYNNMDLTSL---------GGQGH-VGFKV**C**---NQGQD-N**C**VIYTF--------------NS |
| A. Lungfish γ | TRMYNNMDLATV---------GEQ---VGFKI**C**---DANKS-N**C**IIYTF--------------NS |
| Coelacanth γ | RKMFENVAIENQ---------GEL---VGFKL**C**---DTNGS-D**C**AIYTF--------------NS |
| Frog γ | HRDAESV--------------QDPGNMVGFKL**C**---DPKNSSD**C**TIFTF--------------SS |
| Cow γ | HKASDVMHI------------HNSKEVVGFQL**C**---SNDTS-D**C**AVYTF--------------SS |
| Human γ | HKASNVMHI-------------ESKQVVGFQL**C**---SNDTS-D**C**ATYTF--------------SS |
| Chicken γ | HKDSSIVNS------------GDSNDIIGFQL**C**---DANNSSE**C**ALYTF--------------SS |
| Turtle γ | HQGSSMVNT------------GDPQDVVGFQL**C**---DPNNSSD**C**AVYTF--------------SS |

```
Spotted Gar γ-like      ----IEDSNQL------GF--LLSKLN-----PDQQAEAGHQLEDMLISCHFHGEKCD-K-----
Asian Arowana γ-like    -----------------KFASEFNRLS-----DEEKLDMGHQLEDMLLFCNYHGQPCN-T-----
Coelacanth ASIC1        PAQ-----YQL------NTTDLINRL-------------GHQMEEMLLECRFQGETCT-S-----
Catfish ASIC1           PPR-----FHL------NTTEMIGRLS-------------HQLEDMLLLCRFRGESCT-Y-----
E. Lamprey ASIC1        RPFSMREFYE---------------------------RAGHELREMLLHCKFHGMNCT-P-----
E. Shark ASIC1          KPFNMREFYA---------------------------RAGHDMKDMLLHCIFKGEFCT-A-----
J. Medaka ASIC1         RSFNMREFYD---------------------------RTGHDIKDMLLSCSYRGTECS-A-----
Black Rock Cod ASIC1    RPFNMREFYD---------------------------RTGHDIKEMLLSCSYRGVECS-A-----
Lancelet γ-like         DFEMVN-----------SFMGLLVQLT-----ASMRQSLGHQGRDFIQECQFDGRTCS-H-----
Lancelet α-like         SSDYMSEDYHHEFELVQNFSSSILGLN-----RTSRRTMGHQYQDLVLECAYDGRSCS-R-----
E. Shark α              G----------------------------------------RSSEFIRSCKFNRVSCD-N-----
Frog δ                  GVNALHEWYKF------HYINIMSNIP------AVLNIANNFSKDFILTCHFNEVPCD-E-----
Coelacanth α            GMDAVREWYMF------HYVNIMVQVP-----MVTNPLQETHIRDFVFSCKFNHASCN-Q-----
S. Lamprey α            GVQAVTEWYTF------QYVNIMSQVP-----SYIKQSDDANIEDFIFSCMFSGMPCS-D-----
J. Lamprey α            GVQAVTEWYTF------QYVNIMSQVP-----SYIKQSDDANIGDFIFSCMFSGMPCS-D-----
W. Lungfish α           GVDALREWYKY------QYVNIMGNAP-----LSTYQEDNPQISNFVYACEFNKISCG-S-----
A. Lungfish α           GVDALREWYKF------QYINIMAQIP-----SQSNQEDDSQISNFVYACEFNKVSCG-V-----
Ropefish α              GVDAIREWYKY------HYINIMAQMSMNALADDDSNPDKADINNFVFACSFNGAVCS-K-----
Frog α                  GVDAIREWYRF------HYINILARVP------QEAAIDGEQLENFIFACRFNEESCT-K-----
Salamander α            GVDAIREWYRF------HYINILARVP----ATSGVPLNEDSFQNFIFACRFNEDSCS-E-----
Cow α                   GVDAVREWYRF------HYINILSRRR----QDTSPSLEEDVLGKFIFTCRFNQDSCN-E-----
Human α                 GVDAVREWYRF------HYINILSRLP-----ETLPSLEEDTLGNFIFACRFNQVSCN-Q-----
Chicken α               GVDAVREWYSF------HYINILAQMP------DAKDLDESDFENFIYACRFNEATCD-K-----
Turtle α                GVDAVREWYSF------HYINILARMP------NTKALDESNFENFIYACRFNEVTCD-K-----
Cow δ                   GVVAAREWYRF------HYINILALLP--------AAHEDSHGSHFVFSCRYDDRDCH-A-----
Human δ                 GVAAVQDWYHF------HYVDILALLP-----AAWEDSHGSQDGHFVLSCSYDGLDCQ-A-----
Coelacanth δ            VMDAIQEWYKF------HFINIMSQVS-----PMTNVSDSSPIGNVIYSCQYNGKSCS-G-----
Chicken δ               GMDAILEWYRF------HYMNIMSQQP----VIINISDHEEKIEDMVYSCQYDGEPCR-P-----
Turtle δ                GVDTIQEWYRF------HYMNIMSQLP----VIINISSHEEHIQNLVYSCQYDGEPCR-E-----
S. Lamprey β            GMTAVKQWFHF------NLLSLLGNLS-----TEEKSNLSSSGDELIRSCLFSSDTCS-A-----
J. Lamprey β            GMTAVKQWFHF------NLLSLLGELS-----NEEKSNLSSSGDELIRSCLFSNDACN-A-----
E. Shark β              GLQAVNEWYSL------HYMSLMANVS-----LEDRTAMGEHGQDFILSCNFGGHPCD-L-----
Coelacanth β            FLSRVTHNYVQ------NFFFT----------DSSLVYAPTEGRLSFLQVHYG-EMCA-YFSWTL
Ropefish β              GVDAVNEWYSL------HFMDIMSKFS-----INEKKQMAYSGKEFILTCLFGNQPCS-Y-----
W. Lungfish β           SLEAVKEWYTL------HYLNIMLRIP-----LAEKAAMGYSGKDLILTCFFGGIACD-Y-----
A. Lungfish β           SLEAVKEWYTL------QYIDIISKLP-----LSQKVEMGYSGKDFILTCLFGGEACN-Y-----
Frog β                  GVQALREWYLL------QLSSIFSNVP-----LSGRIDMGFKAEDLILTCLFGGQPCS-Y-----
Cow β                   ATQAVTEWYTL------QATNIFAQVP-----NQELVAMGYPAERLILACLFGAEPCN-Y-----
Human β                 ATQALTEWYIL------QATNIFAQVP-----QQELVEMSYPGEQMILACLFGAEPCN-Y-----
Chicken β               AAQAVTEWYIL------QSTSILSKVP-----LQERIRMGYQAEDMILACLYGAEPCN-Y-----
Turtle β                AAQAVTEWYIL------QSTSILSKVP-----LQERIRMGYQPEDMILACLYGAEPCS-Y-----
S. Lamprey γ            AIDAVIQWYRL------HFINIMAIVP-----EKDKDKLGYSADEFIIDCLFSGTVCDPS-----
J. Lamprey γ            AIDAVMQWYRL------HFINIMAIVP-----EKDKDKLGYSADEFIIGCLFSGTVCDPS-----
Ropefish γ              GISAVQEWYWL------HFNNIIAQQS-----LETLVEMGYSAEEFISTCTFNEAMCS-L-----
E. Shark γ              GMAAVQEWYQL------HYYNLLAQVP-----PEDKRAMGYSADDLFLTCLYDGLPCD-S-----
W. Lungfish γ           GITALQEWYRL------NFIDIMAQVP-----NEKKAEMGYSADELIVSCMYDGQACD-S-----
A. Lungfish γ           GVTAILEWYRL------NYLNIMAQIP-----NEKKLEMGYSADDLIVTCMYDGQSCD-S-----
Coelacanth γ            GITAIQEWYRL------HYINIMAQVS-----WEKKQEMGYSADDLIVTCFYNGMPCN-S-----
Frog γ                  GVNAIQEWYRL------HYTNILAKIS-----MEDKIAMGYKADELIVTCFFDGLSCD-A-----
Cow γ                   GVNAIQEWYKL------HYMNIMAQVS-----QEKKINMSYSADELLVTCFFDGVSCD-A-----
Human γ                 GINAIQEWYKL------HYMNIMAQVP-----LEKKINMSYSAEELLVTCFFDGVSCD-A-----
Chicken γ               GVNAIQEWYKL------HYMNIMAQIP-----LETKEELSYSADDLLLTCFFDGLSCD-K-----
Turtle γ                GVNAIQEWYKL------HYMNIMAQIP-----LETKVNMSYSAEDLLLTCFFDGISCD-T-----
```

```
Spotted Gar γ-like       ---SFFNAFFNHKFGNCYTFNSLTKMENRGRLMR-RDVLNATKAGFSYGLTMELSIEQDEYIEQF
Asian Arowana γ-like     ---SFFSGFINYKFGNCYTFNSHKQTDIRGRPIK-SESLNTTKAGFMYGLHLELFIQQNEYVRDI
Coelacanth ASIC1         ---RNFTPIF-TRYGKCYTFNS-------G--KDGNPLLTTLKGGMGNGLEIMLDIQQDDYLPVW
Catfish ASIC1            ---KNFTTIY-TRYGKCYTFNS-------G--LDGNPLLTTLKGGTGNGLEIMLDIQQDEYLPVW
E. Lamprey ASIC1         ---QDFQTVY-TRYGKCYTFNS-------G--KDGRPLLTSMKGGMGNGLEMMLDIQQDEYLPVW
E. Shark ASIC1           ---QDFKIVF-TRYGKCYTFNS-------GQIKD-QPILTTLEGGTGNGLELMLDIQQDEYLPVW
J. Medaka ASIC1          ---ENFKVIF-TRYGKCYTFNS-------G--KDGQPLMVTMKGGTGNGLELMLDIQQDEYLPVW
Black Rock Cod ASIC1     ---DNFKVIF-TRYGKCYTFNS-------G--QDGRPLMVTMKGGMGNGLELMLDIQQDEYLPVW
Lancelet γ-like          ---RNFTTFEDSTYGNCFTFNK-------D--KDGEVLHTATSAGPLHGLSLILYIEQDEYIPAI
Lancelet α-like          ---TDFGRILDEKYGNCYTFNS-------D--K--VLQRKVRDPGPAHGLQLTLYIEQDEYVPAV
E. Shark α               ---NNYTTFSHPRYGTCYTLNT-------G-----PVSWRVLGPGSANGLTLTLQVGEG--LRFL
Frog δ                   ---REYIHFHHPIYGNCFTINN-------H--GK-ENSWYSPRPGKQYGLSMVVKADLHDNMPLL
Coelacanth α             ---GNYTYFNHPVYGNCYTFNG-------G--ST-GNLWSSTKPGRENGLSLLLRTEQNDYIPFL
S. Lamprey α             ---SEYSRFHHPTYGNCYTFNS-------A--NS-SKLWQASKPGRDYGLSLILRTEQNDYIPFL
J. Lamprey α             ---SEYSRFHHPTYGNCYTFNS-------A--NS-SKLWQASKPGRDYGLSLILRTEQNDYIPFL
W. Lungfish α            ---GNYTQFNHPQYGSCYTFND-------G--DD-NNPWISFSPGVESGLSLVLRTEQNDFLPYL
A. Lungfish α            ---ENYTRFRHPVYGNCYTYND-------G--QS-ATPWASFVPGVGNGLSLVLRTEQNDFLPFL
Ropefish α               ---GNYTTFHHPMYGNCYTFNS-------W--ED-GHEWSVSTPGVESGLSLLLRTEQNDFIPLL
Frog α                   ---ANYSSFHHAIYGNCYTFNQ-------NQSDQ-SNLWSSSMPGIKNGLTLVLRTEQHDYIPLL
Salamander α             ---ANYTHSHHPLYGNCYTFNE-------DHSRN-DSRWASSMPGINYGLSLVLRTEQNDYIPLL
Cow α                    ---ANYSHFHHPMYGNCYTFND-------K--NS-SNLWMSSMPGVNNGLSLTLRTEQNDFIPLL
Human α                  ---ANYSHFHHPMYGNCYTFND-------K--NN-SNLWMSSMPGINNGLSLMLRAEQNDFIPLL
Chicken α                ---ANYTHFHHPLYGNCYTFND----------NS-SSLWTSSLPGINNGLSLVVRTEQNDFIPLL
Turtle α                 ---ANYTHFHHPIYGNCYTFND----------GN-SSLWTSSLPGINNGLSLLVRTEQNDYIPLL
Cow δ                    ---QHFQTSHHPTYGSCYTFNG---------------VWAAQRPGVTHRISLVLRAEQLDHLPLL
Human δ                  ---RQFRTFHHPTYGSCYTVDG---------------VWTAQRPGITHGVGLVLRVEQQPHLPLL
Coelacanth δ             ---SEYEHFHHPVYGICYIFKS-------N--GS-DTFWETSKPGIAYGLSLIIGTKQEDFIPLL
Chicken δ                ---SDYVHFHHPVFGSCYTFNS-------K--GT-DPFWTATKPGIPYGLSLILRAEQKDHIPLL
Turtle δ                 ---SDYIHFHHQVYGSCYTFNS-------E--GT-DLFWKASKPGISYGLSLILKAEQNDRLPLL
S. Lamprey β             ---TNFTTFFHPMYGNCFIFNW-------G--EN-ETVMQVSNPGVEYGLKLVLSIDQDEYIPFL
J. Lamprey β             ---TNFTTLYHPMYGNCFIFNW-------G--EN-ETVMQVSNPGVEYGLKLVLSIDQDEYIPFL
E. Shark β               ---RNFTRLFHPTYGNCYIFNW-------G--SS-GSVLTVSNPGAEFGLKVVLDISQEDYNPFL
Coelacanth β             LTVINFTQMFHPTYGNCYIFNW-------G--QD-GNALISSNPGADFGLKLVLDINQEEYIPFL
Ropefish β               ---KNFSQIFHHFYGNCYIFNW-------G--LH-DKAISSSNPGGEFGLNVVLDINQKEYIPFL
W. Lungfish β            ---TNFTQIYHPSYGNCYIFNW-------G--LD-GNVAVSSNPGVGFGLQLVVDVNQEEYIPFL
A. Lungfish β            ---DNFTQFYHSSYGNCYVFNW-------G--LD-GNVLIVSNPGVGFGLQLALDVNQEEYIPFL
Frog β                   ---RNFTHIYDADYGNCYIFNW-------G--QEGENTMSSANPGADFGLKLVLDIEQGEYLPFL
Cow β                    ---RNFTPIFHPDYGNCYIFNW-------G--MT-EKALPSANPGTEFGLKLILDMGQEDYVPFL
Human β                  ---RNFTSIFYPHYGNCYIFNW-------G--MT-EKALPSANPGTEFGLKLILDIGQEDYVPFL
Chicken β                ---KNFTQIYHPDHGNCYIFNW-------G--MD-KEALNSSNPGAEFGLKLILDISQQDYIPYL
Turtle β                 ---RNFTQIYHPDHGNCYIFNW-------G--MD-EEALISSNPGAEFGLKLILDISQQDYIPYL
S. Lamprey γ             ---TSFKKLQHPILGNCFTFND-------G--SD-GKSLDIASAGIDYGLHMVLNTRQDNSLPYL
J. Lamprey γ             ---TSFKKLQHPILGNCFTFND-------G--RD-GKSLDIASAGIDYGLHMILNTRQDNALPYL
Ropefish γ               ---RNFTQSYHPTLGNCYTFNS-------G--YD-GEIIQSSTAGIKNGLIVVLNLGLEDYNPFL
E. Shark γ               ---RNFSLHQHPLHGNCFTFNG-------G--EN-GRVLITRTGGSQNGLKVTLHLDEDEYNPYL
W. Lungfish γ            ---RNFTLFQHPLHGNCYTFNS-------G--ND-GNILQTLIGGNARGLKLILYTENDDYNPFL
A. Lungfish γ            ---RNFTLFQHPLHGNCYTFNS-------G--DD-GNILQTLTGGSEYGLKLTLYLENDDYNPYL
Coelacanth γ             ---RNFTLFQHPVYGNCYTFNS-------G--AD-GSILKTSTEASEFGLNVILYIDHKDYNPFL
Frog γ                   ---RNFTLFHHPLYGNCYTFNS-------A--ER-GNLLVSSMGGAEYGLKVVLYIDEDEYNPYL
Cow γ                    ---RNFTLFHHPMYGNCYTFNN-------R--QN-ETILSTSMGGSEFGLQVILYINEEEYNPFL
Human γ                  ---RNFTLFHHPMHGNCYTFNN-------R--EN-ETILSTSMGGSEYGLQVILYINEEEYNPFL
Chicken γ                ---RHFTRFHHPLHGNCYTFNS-------G--EN-GTVLSTSTGGSEYGLQVVLYIDEADYNPFL
Turtle γ                 ---RNFTPFHHPLHGNCYTFNS-------G--EN-GKVLTTSTGGSEYGLQVVLYIEEADYNPFL
```

| | |
|---|---|
| Spotted Gar γ-like | SQA------AGIRLIIHDQKDMPFPEDDGVNIPPGQESDIAIVKVHVHRLRAPYSSTCTTGDGIH |
| Asian Arowana γ-like | THS------AGIRMLIHDHLATPFPEDEGVNIPPGTETDIGITKVGIRRLKHPYGSNCTDGEGIT |
| Coelacanth ASIC1 | KETDETSLEAGIKVQIHSQEEPPFIDQLGFGVAPGFQTFVSCQQQKLMYLPPPWGDCKATPINSE |
| Catfish ASIC1 | GDTDETSYEAGIKVQIHSQDEPPFIDQLGFGVAPGFQTFVSCQQQLLQYLPPPWGDCQSTAMNSE |
| E. Lamprey ASIC1 | GETDETSFEAGIRVQIHSQDEPPFIDQLGFGVAPGFQTFVSCQEQRLTYLPYPWGDCKDTPPESE |
| E. Shark ASIC1 | GETDETSFEAGIKVQIHSQSEPPFIDQLGFGVPPGFQTFVACQEQRLRYLPPPWGDCKSTPMDSD |
| J. Medaka ASIC1 | GETDETSFEAGIKVQIHTQEEPPFIDQLGFGVAPGFQTFVSCQEQRLTYLPPPWGDCKSTPMDSD |
| Black Rock Cod ASIC1 | GETDETSFEAGIKVQIHTQDEPPFIDQLGFGVAPGFQTFVSCQEQRLTYLPPPWGDCKASAMDSD |
| Lancelet γ-like | AEK------AGARVVIHNPYVFPFPESEGFDAAPGFLTSAGLRLTSITRLGGVYGNCT-NGQG-- |
| Lancelet α-like | TPA------AGVRVVIHQPGEWPFPAEEGFDVGPGYSTSIGLQVTTIRRLGGKYGNCT-DGRDKD |
| E. Shark α | TPG------FGVRLMVHDPKQTPILEDDGIDLLPGLETSVSLRLESVRRLGGGLSDCTKDGKGVE |
| Frog δ | SQA------AGARIMIHNPNQPPLVEHEGFDIQPGTETSISVKQEEVIRLGGKYSQCTSDGSDLS |
| Coelacanth α | STV------AGARVMIHRQNQPPFMEDEGFNIRPGVETSISMKKVSRQQLGGLYSDCTEDGSDIG |
| S. Lamprey α | STV------AGARIMVHDQESPPFMEEGGFDMRPGFETSLGIRMLEATRMPDPYGNCTEDGSNVP |
| J. Lamprey α | STV------AGARIMVHDQESPPFMEEGGFDMRPGIETSLGIRMLEATRMPDPYGNCTEDGSDVP |
| W. Lungfish α | SNV------AGARVMVHDQNQPVFMEDSGFDIRPGVETSIGIKKEIISRLGGVYGNCTADGSDIN |
| A. Lungfish α | STV------AGARVLVHDQNQPPFMEDSGLDIRPGVETSIGMKKEIISRLGGVYGNCT-DGSDID |
| Ropefish α | STV------AGARVMVHSQNHPVFMEDGGFDIKPGVETSIGLRQEVFQRLGGEYGDCL-DGTDLD |
| Frog α | SSV------AGARVLVHGHKEPAFMDDNGFNIPPGMETSIGMKKETINRLGGKYSDCSEDGSDVD |
| Salamander α | STT------AGARVIIQAPDEPVLLNEGGFNIQPGVETSISMTKETMDRLGGAYSDCTEDGSDVE |
| Cow α | STV------TGARVMVHERDEPAFMDDAGFNLRPGVETSISMSKEAVDRLGGDYGDCTKNGSEVP |
| Human α | STV------TGARVMVHGQDEPAFMDDGGFNLRPGVETSISMRKETLDRLGGDYGDCTKNGSDVP |
| Chicken α | STV------TGARVMVHDQNEPAFMDDGGFNVRPGIETSISMRKEMTERLGGSYSDCTEDGSDVP |
| Turtle α | STV------TGARVMVHEQNEPAFMDDGGFNVRPGMETSISMRKETTMRLGGSYSDCTEDGSDVL |
| Cow δ | STK------AGIKVMIHPQDHTPFLEHQGFSIRPGTETTIDIREDEVHRLGSPYGQCMDSTGSVD |
| Human δ | STL------AGIRVMVHGRNHTPFLGHHSFSVRPGTEATISIREDEVHRLGSPYGHCTAGGEGVE |
| Coelacanth δ | STV------AGTRVMIHKQDQPAFMEDEGLNIKPGTETSIGMKQDEVNRLAGNYGQCTFDGTDVK |
| Chicken δ | STV------AGVKVMIHNHNQTPFLEHEGFDIRPGIATTIGIQQDKVNRLGGNYGKCTTDGSDVK |
| Turtle δ | STV------AGVQVMIHNHNQTPFLEHEWFDIRPGIATNIGIRQDEVHRLGGNYGKCTVDGADVD |
| S. Lamprey β | TTI------AGAVIMVHDQNTYPFLSDLGVFVKTGVETSVGIEVGQLQRQGAPYSDCTMDGTDLP |
| J. Lamprey β | TTI------AGAVIMLHDQNTYPFLSNQGFFVKTGAETSVGIEVGQLQRQGAPYSDCTMDGTDLP |
| E. Shark β | SMA------AGAKFMLHQQNTFPFLRDLGMYAKAGTESSITIFADEIERLGGVYSRCRLNPSATE |
| Coelacanth β | TTS------AGARLMLHDQNTFPFLKDLGMYAMAGSQTSIGILVDEIQRIGAPYSQCTPYGSDVP |
| Ropefish β | STT------AGARLLVHEQRSFPFLKDLGIFVLPGTETSIGISVDKIERMEAPYSDCTQNGSDVP |
| W. Lungfish β | TTS------AGVRFLLHDQKTFPFVETMGIYALVGTVTSVEILVDEVMRMEQPYGTCTADGSDVP |
| A. Lungfish β | TTR------AGARFLLHTQNTFPFVETMGTYALVGTVTSVGILVDEVQRMGQPYGTCTTDGLDVP |
| Frog β | QTT------AAARLILHQQRSFPFVKDLGIYAMPGTETSISVLVDQLEHMEAPYSSCTVNGSDIP |
| Cow β | TST------AGARLMLHEQRSYPFIKEEGIYAMAGMETSIGVLVDKLQRKGEPYSQCTKNGSDVP |
| Human β | AST------AGVRLMLHEQRSYPFIRDEGIYAMSGTETSIGVLVDKLQRMGEPYSPCTVNGSEVP |
| Chicken β | SSA------AGARLMLHQQKSFPFLKDQGIYAMAGTETSIGVLVDELERMGYPYSDCTANGSDVP |
| Turtle β | TST------AGARLMLHEQKSFPFLKDQGIYAMSGTETSIGVLVDELERMGYPYSDCTMNGSDVP |
| S. Lamprey γ | AMG------AGAKIGIHLQNTTPFIEAVGIDIPPAMESSLGLRVNDVQKLGDPYSDCTMDGSDID |
| J. Lamprey γ | AMG------AGAKIGIHLQNTTPFIEAVGINIPPATESSLGLRVNDVQKLGEPYSDCTMDGSDID |
| Ropefish γ | SSS------EGAIIMIHNQNEHPFIEDLGIMIQTAKETSIGLQFMESHKLGEPYSSCTEDGTDVS |
| E. Shark γ | VTS------TGAKIVVHDQSEHPFVEDLGIAIPAGMETSIGLDLTESHKLGGPYSDCIE---DLP |
| W. Lungfish γ | FTS------MGAKVIVTDQNEYPLIDDVGLEVQTAMETLVGLQLTDSAKLSQPYSDCTVDGSDVM |
| A. Lungfish γ | FTS------MGAKIIVHDQTEYPLVDDVGLEIQTATETLIGLQVTTSAKLSKPYSDCTMDGSDVL |
| Coelacanth γ | VTS------TGAKVVIHDQNEHPFIEDMGLEVETATETSIGLQLTESHRLSSPYSNCTEDGSDVP |
| Frog γ | STA------AGAKILVHDQDEYPFIEYLGTELETATETSIGMQLTESAKLSDPYSDCTMDGRDVS |
| Cow γ | VSS------TGAKVIIHRQDEYPFVEDVGTEIETAMATSIGMHLTESFKLSDPYSQCTEDWSDVQ |
| Human γ | VSS------TGAKVIIHRQDEYPFVEDVGTEIETAMVTSIGMHLTESFKLSEPYSQCTEDGSDVP |
| Chicken γ | VTS------TGAKIIVHDQDEYPFIEDIGTEIETAAATSIGMHFTRSRKLSKPYSDCTETGADIP |
| Turtle γ | VTS------TGAKIIVHDQNEYPFIEDIGTEIETATATSIGMHFTRSHKLSKPYSDCTETGTDIP |

```
Spotted Gar γ-like      NYYR--DVYK-VGYSR-------EACKKTCGQMYIIKNCGCGMWEFPVPKDVKVPFCNITNKNI-
Asian Arowana γ-like    NFYH--DLHG-FKYTR-------EACKRTCAQQSIMKDCGCSHWEFAVLPDLQYPKCNFSSPAT-
Coelacanth ASIC1        --------FF-STYSI-------TACRIDCETRYLVENCNCK--MVHMPGNAKV--CTPDQY---
Catfish ASIC1           --------FF-STYSI-------TGCRIDCETRYLLENCNCR--MVHMPGTSTV--CTPEQY---
E. Lamprey ASIC1        --------FF-DTYSI-------AACQIDCETRYLVENCNCR--MVHMPGDAPY--CTPEQY---
E. Shark ASIC1          --------FF-DTYSI-------TACRIDCETRYLVENCNCR--MVHMPGDAPY--CTPEQY---
J. Medaka ASIC1         --------FF-NSYSI-------TACRIDCETRYLVENCNCR--MVHMPGDAPY--CSPEQY---
Black Rock Cod ASIC1    --------FF-NTYSI-------TACRIDCETRYLVENCNCR--MVHMPGDAPY--CTPEQY---
Lancelet γ-like         RHLL----YP-QMYSQ-------QNCLATCHQEHMVEICGCADVTFLQPN---------------
Lancelet α-like         --NL----YR-SKYST-------KTCLHTCFQRLLVEKCGCGSRFIPLPKKVPA--CPVPI----
E. Shark α              IENL----YD-SSYSQ-------QTCVRSCFQALMTLRCNCSYFFYNKPKNSHY--CNSRSHPDW
Frog δ                  IKIL----YN-TSYTM-------QACLNSCFQYKMIEMCGCGYYFYPLPPGMEY--CNYNKYPGW
Coelacanth α            VENL----YN-SNYTQ-------QACVRSCFQVTLVQRCGCGHYFYPLPEGAQY--CNYKKHKTW
S. Lamprey α            VLNL----YS-SAYTV-------QVPECSCFQLALVEACGCGYYFYPLPPNASY--CSYNN-TAW
J. Lamprey α            VLNL----YS-SAYTV-------QACVRSCFQLALVEMCGCGHYFYPLPPNAAY--CSYNN-TAW
W. Lungfish α           VENL----YN-SDYTQ-------QACIRSCFQATIVERCGCGYYFYPLPAGATY--CTNTKHRGW
A. Lungfish α           VVNL----YN-SDYNQ-------QACVRSCFQATIVQQCGCGYYFYPLPSGAEY--CSYSRNKSW
Ropefish α              IENL----YE-SSYTQ-------QACIRSCFQLIMVKRCGCAYYFYPLPKGASY--CNYNRHIAW
Frog α                  VKNL----FQ-SEYTE-------QVCVRSCFQAAMVARCGCGYAFYPLSPGDQY--CDYNKHKSW
Salamander α            VKNL----FN-KKYTQ-------QACVRSCFQANMVQRCGCAYYFDPLPPGEEY--CDYHKQPNW
Cow α                   VENL----YN-TKYTQ-------QVCIHSCFQESMIKECGCAYIFYPRPDGVEF--CDYRKHNSW
Human α                 VENL----YP-SKYTQ-------QVCIHSCFQESMIKECGCAYIFYPRPQNVEY--CDYRKHSSW
Chicken α               VQNL----YS-SRYTE-------QVCIRSCFQLNMVKRCSCAYYFYPLPDGAEY--CDYTKHVAW
Turtle α                VQNL----YS-SRYTE-------QVCIRSCFQSSMVERCGCAYYFYPLPSGAEY--CDYTKHIAW
Cow δ                   VQLL----YN-TSYTR-------QACLVSCFQHLMVETCSCGYFFYPLPAGAEY--CSYMRHPAW
Human δ                 VELL----HN-TSYTR-------QACLVSCFQQLMVETCSCGYYLHPLPAGAEY--CSSARHPAW
Coelacanth δ            IK-L----YN-TPYSV-------QACVRSCFQYLLIQECGCGYYYYPLPPGAQY--CNYNKYPSW
Chicken δ               VKLL----YN--SYTL-------QACLHSCFQHIMVQKCGCGYYYYPLPPGAEY--CNYNKQPAW
Turtle δ                VKLL----YN-SSYTL-------QACLHSCFQDKMVERCGCGYYYYPLPPGAEY--CNYNKHPAW
S. Lamprey β            ITNL----YNGTAYSV-------QACLRSCFQTKMIEMCGCGYYLYPLPPGEKY--CQNQNFTGW
J. Lamprey β            ITNL----YNGTAYSV-------QACLRSCFQTKMIEMCGCGYYLYPLPPGEKY--CQNQNFTGW
E. Shark β              VTTL----YN-TSYSM-------QTCLRSCHQAHMVRLCGCAYHHYPLSEGAQY--CNNQDHPGW
Coelacanth β            VPNL-YSIYN-TSYSM-------QNCLYSCLQAKLVEKCGCGNYLHPLPDGAHS--CNNEDNPSW
Ropefish β              ITDLFYKMYE-TSYSV-------QSCLRSCFQMNLVKMCGCAYNLYPLPAEAPY--CNFGDHPEW
W. Lungfish β           VNNL-YSSYN-LSYSM-------QSCLWSCFQAQMVKRCGCAYYLYPLLDGANY--CDTQNNSDW
A. Lungfish β           IDNL-YSQYN-LSYTM-------QSCLWSCFQIQMVNSCGCAYYLYPLPEGATY--CNNQNNSDW
Frog β                  VQNL-YAEFN-SSYSI-------QSCLRSCYQEEMVKTCKCAHYQYPLPNGSEY--CTNMKHPDW
Cow β                   IQNL-YSNYN-TTYSI-------QACIRSCFQEHMIRECGCGHYLYPLPHKRKY--CNNQEFPDW
Human β                 VQNF-YSDYN-TTYSI-------QACLRSCFQDHMIRNCNCGHYLYPLPRGEKY--CNNRDFPDW
Chicken β               VKNL-YSEYN-TSYSIQLPLSFFQACLRSCFQNHMTEICGCGHYMFPLPEGVTY--CNNEDNPGW
Turtle β                VKNL-YNEYN-TSYSI-------QACLRSCFQAQMFENCGCGHYLFPLPEGVNY--CNNEDDPDW
S. Lamprey γ            VKSL----YD-SPYSV-------QTCQNSCFQWEMIKSCGCANYEQPLPEGSRF--CNYDNNPGW
J. Lamprey γ            VKSL----YD-SPYSV-------QTCQNSCFQLEMIKSCGCANYEQPLPEGSLF--CNYDNNPGW
Ropefish γ              VNNL----YN-KTYSL-------QVCLHSCFQKEMTLQCGCAHFHYPLPAEAQY--CNYNAFPDW
E. Shark γ              EDNL----YN-KSYSL-------QMCLHSCFQKEMVQTCGCGHYEKPLPPGAQY--CDYNRFPGW
W. Lungfish γ           EENL----YN-KSYSL-------QICLHSCFQKEMVNSCGCAYYEQPLPPGAEY--CSYEKFPGW
A. Lungfish γ           EQNL----YN-TSYSL-------QICLHSCFQTEMISNCGCAYYEQPLPSGAEY--CYYEKYPGW
Coelacanth γ            MQNL----YN-TTYSF-------QMCLYSCFQKEMVQSCGCAHYEYPLPVDTEY--CDYKKYPGW
Frog γ                  VENL----YN-KKYTL-------QICLNSCFQREMVRSCGCAHYDQPLPNGAKY--CNYEEYPSW
Cow γ                   ITNI----YN-ATYSL-------QICLHSCFQAKMVENCGCAQYSQPLPRGADY--CNYQQHPNW
Human γ                 IRNI----YN-AAYSL-------QICLHSCFQTKMVEKCGCAQYSQPLPPAANY--CNYQQHPNW
Chicken γ               VENL----YN-KSYSL-------QICLHSCFQKAMVESCGCAQYAQPLPNGAEY--CNYKKNPNW
Turtle γ                VANL----YN-KSYSL-------QICLHSCFQRAMVDTCGCAQYAQPLPPGAEY--CNYKKYPNW
```

| | |
|---|---|
| Spotted Gar γ-like | --NKCVQLYEDKFAHDELEC--NCPLQCEEEIFELTLSSSQWPSAVYMNEFARKLRQSG------ |
| Asian Arowana γ-like | --RRCLELYEYKFAQDILPC--HCPLQCKEELYSLTVSGSQWPATAFLDKFSSNLRGKG------ |
| Coelacanth ASIC1 | --KNCADPALDFLVEKD-NDYCVCQTPCNMTRYGKELSMVKIPSKASAKYLAKKFNKTE------ |
| Catfish ASIC1 | --KDCADPALDFLVEKD-NNYCVCPTPCNMTRYGKELSMVKIPSKASAKYLAKKFNKSE------ |
| E. Lamprey ASIC1 | --KECANPALMFLVEKD-DDYCACEMPCNIKRYAKELSMVKVPSQASAKYLAKKFNKTE------ |
| E. Shark ASIC1 | --KECADPALDFLVEKD-SVFCTCETPCNMTRYGKELSMVKIPSKASAKYLAKKYNKSE------ |
| J. Medaka ASIC1 | --KECADPALDFLVERD-NDYCVCETPCNLTRYGKELSFVKIPSKASAKYLAKKFNKTE------ |
| Black Rock Cod ASIC1 | --KDCADPALDFLVERD-NDYCVCETPCNMTRFGKEMSFVKIPSKASAKYLAKKFNKTE------ |
| Lancelet γ-like | --VDCEEKVKQRLGNGNLTC--QCPISCMDRIYRKAIGLSEWPADSYVSTVLNKLKTKR------ |
| Lancelet α-like | --NSCEQRWISEMRNGRIWC--DCPPSCVDNTLSMTFGFSEWPADSYEKSLRQKLSKLD------ |
| E. Shark α | --GHCYYKLYEEFIAEKLNCFEKCPKLCDQSLYHITVGHSKWPSKQVESWMRPFLSNKK------ |
| Frog δ | --GHCFYQLYEKMLDHTLICFTQCPKQCKQTQYHLAAGTAKWPSPVSKA--IQLLSLQE------ |
| Coelacanth α | --GHCYYRLYKEFKANDLGCFTKCRKRCLESEYHQMTGYSKWPAKDSGKWIHHILAKQN------ |
| S. Lamprey α | -AGHCYYKLYRQFISDELGCVDKCAQPCTTKRFAVTPGYAAWPDSSSEKWIFNLLSLQN------ |
| J. Lamprey α | -AGHCYYKLYRRFISDELGCVDKCAQPCTTKRFAVTPGYATWPDSSSEKWIFNLLSLQN------ |
| W. Lungfish α | --GYCYYKLYKAFAADELGCFKRCPKPCIVAEYVKTAGYSKWPSSSSETWIAKVLSQES------ |
| A. Lungfish α | --GYCYYKLYKAFAADELGCFRRCRKPCQYTDYKMTAGYAQWPSSVSESWITSILSQEN------ |
| Ropefish α | --GHCYYKLYNEFSLDNLGCSTKCRRPCQDTEYTMTAGYATWPTKASKNWIFNVLNKQN------ |
| Frog α | --GHCYYKLIIEFTSNKLGCFTKCRKPCLVSEYQLTAGYSKWPNRVSQDWVLHTLSR-------- |
| Salamander α | --GHCYYKLENEFVSDDLQCFTKCRKPCQLSEYHLSAGYSRWPSDVSKSWVFHMLSQQN------ |
| Cow α | --GYCYYKLQDAFSSDRLGCFTKCRKPCSVTIYKLSASYSQWPSATSQDWVFQMLSRQN------ |
| Human α | --GYCYYKLQVDFSSDHLGCFTKCRKPCSVTSYQLSAGYSRWPSVTSQEWVFQMLSRQN------ |
| Chicken α | --GYCYYKLLAEFKADVLGCFHKCRKPCKMTEYQLSAGYSRWPSAVSEDWVFYMLSQQN------ |
| Turtle α | --GHCYYKLQVEFKSNVLGCFSKCRKPCEVTEYQLSAGHSHWPSTVSEDWVSHMLSRQN------ |
| Cow δ | --GHCFHHLYQKLKTHQLPCTTRCPRPCRESSYKLSAGTSRWPSSTSADWVLAVLGEPSRRNPWP |
| Human δ | --GHCFYRLYQDLETHRLPCTSRCPRPCRESAFKLSTGTSRWPSAKSAGWTLATLGEQG------ |
| Coelacanth δ | ETGHCYYKLYKKFVAGDSGCFQKCPKPCQEFKYKLTTGISKWPSQNAENWIFHLLSHHN------ |
| Chicken δ | --GHCFYQLYSRLRNHHLNCFDQCPKPCRESLYKVSAGTAKWPSRKSQDWIRQALRHQN------ |
| Turtle δ | --GHCFYQLYNRLADHHLSCFAKCPKPCWESWYKLSAGTAKWPSTKSQDWVRQILSRQK------ |
| S. Lamprey β | --RYCYYKLYEQFVEEDMDCYTICKQPCIESEYKMSISMSDWPSQSSEDWIFHILSKER------ |
| J. Lamprey β | --RYCYYKLYDQFVEEDMDCYTICKQPCIESEYKMSISMSDWPSQSSEDWIFHILSKER------ |
| E. Shark β | --AYGYYHLKEQIESENSECLTSCIPPCNDTLYRLTISMAEWPSQASEEWIYQILSYER------ |
| Coelacanth β | --AYCYYSLGDSSEYKD-SCLQICEQPCNETQYRLTISMADWPSESSEDWIFHVLSYER------ |
| Ropefish β | --VYCYHKWKKS-AESELLCLQTCKQSCNESQHLLTVSMADWPSESSEDWIFHVLSYER------ |
| W. Lungfish β | --VYCYYHLQDSTDANE-ECIQICELPCIENQFRISTSMADWPSESSEDWIFHVLSHER------ |
| A. Lungfish β | --AYCYYLLQDSKDHKN-ECLQTCIQTCNELQFRISTSMADWPSESSEDWIFHVLSYER------ |
| Frog β | --VPCYYSLRDSVAIRE-NCISLCQQPCNDTHYKMVISMADWPSAGAEDWIFHVLSYEK------ |
| Cow β | --AHCYSALRISLAQRE-TCIYACKESCNDTQYKMTISMAVWPSEASEDWIFHVLSQER------ |
| Human β | --AHCYSDLQMSVAQRE-TCIGMCKESCNDTQYKMTISMADWPSEASEDWIFHVLSQER------ |
| Chicken β | --AYCYSSLRSSIRHRQ-ICIDSCKETCNDTQYKMTISMADWPSEASEDWIFHILSYER------ |
| Turtle β | --AYCYSSLRSSIRHRQ-FCIDSCKETCNDTQYKMTISMADWPSEASEDWIFHILSYER------ |
| S. Lamprey γ | --EYCYYRLYDMYIKEELKCIQVCRQICSETEHEVTLSLADWPSKASKGWLLRALSKEQ------ |
| J. Lamprey γ | --EYCYYRLYDMYIKEELKCIQVCRQICSETEHEVTLSLADWPSKASKGWLLRALSEER------ |
| Ropefish γ | --MVCYSKLHAKFLQEELNCQKTCKGTCHTKEWILTESVAQWPSVNSEKWVLQTLRFGN------ |
| E. Shark γ | --IYCYYRLRDRFHREQLVCQELCRQACHCKEWTLMTSVAQWPAQSAEDWVLRLLSWER------ |
| W. Lungfish γ | --IYCYYQLQDKFVNERLPCQDVCKEPCNSKDFEITKSLAQWPSGASEAWVIRLLDWER------ |
| A. Lungfish γ | --IYCYYQLQDKFVNERLACQDICKETCNSKDWDLTKSLARWPSVASKDWVLNLLNWER------ |
| Coelacanth γ | --IYCYYKLRGKFAQEQLECQQVCKEACNSKEWALTKSLAHWPSLASEDWILRALNLQG------ |
| Frog γ | --IYCYFKVYKQFVQEELGCQSACRESCSFKEWTLTRSLAKWPSLNSEEWMLRVLSWEL------ |
| Cow γ | --MYCYYQLHQAFVREELGCQSVCKEACSFKEWTLTTSLAQWPSEVSEKWLLSILTWDQ------ |
| Human γ | --MYCYYQLHRAFVQEELGCQSVCKEACSFKEWTLTTSLAQWPSVVSEKWLLPVLTWDQ------ |
| Chicken γ | --MYCYYRLHEKFVKEQLGCQQICKDACSFKEWALTTSIAQWPSTVSEDWMLRVLSWDK------ |
| Turtle γ | --MYCYYKLHETFVKEQLGCQQICKEACSFKEWTLTTSLAQWPSSVSEDWMLRVLSWDK------ |

```
Spotted Gar γ-like      ----------GKLAKVADK---VRD--------------------NLVKVIINYQQLNYELIEEI
Asian Arowana γ-like    ----------GQLKAIADNPQDIRD--------------------NMVKVVVYYQKLNYEHISEE
Coelacanth ASIC1        ------------DYIA--------E--------------------NILVLDIFFEALNYETIEQK
Catfish ASIC1           ------------QYIG--------E--------------------NILVLDIFFEALNYEKIEQK
E. Lamprey ASIC1        ------------AYIA--------E--------------------NVLVLDIFFEALNYETIEQK
E. Shark ASIC1          ------------DYIG--------E--------------------NILVLDIFFEALNYETIEQK
J. Medaka ASIC1         ------------QYIA--------D--------------------NILVLDIFFEALNYETIEQK
Black Rock Cod ASIC1    ------------QYIS--------D--------------------NLLVLDIYFEALNYETIEQK
Lancelet γ-like         ---------RGPGTGILDDRDEFKK--------------------NLLKLNIYYEALNYETITES
Lancelet α-like         ----------EESQEKLQTSHDARR--------------------NLLKLSVYFEQLNQQTISES
E. Shark α              ------------DFNV-------RK--------------------DFAKLNVYFEALSYREIEEI
Frog δ                  ------------RYNSTSE----RS--------------------DVSKINVYYEELSYRSVEET
Coelacanth α            ------------QYNFTTN---SRE--------------------DVSKLTVYFQELHHKTVGES
S. Lamprey α            ------------NYSVTTV----RN--------------------DVAKLNVYFRELNMKTISES
J. Lamprey α            ------------NYSVTTV----RN--------------------DVAKLNVYFRELNMKTISES
W. Lungfish α           ------------PYS-TSA----RK--------------------VIAKLNIYFYELSYKTTGES
A. Lungfish α           ------------QYNMTSG----RK--------------------NIAKLNVYFYELNYQTMGES
Ropefish α              ------------GYNITSD----RN--------------------DIAKVNIYFEDLNYRTFGES
Frog α                  ------------QYNLT-D----RN--------------------GIAKLNIYFEELNYKTILES
Salamander α            ------------QYNFTSD----RS--------------------GVAKLNIYFSEMTYKSTVES
Cow α                   ------------NYTIKNK----RD--------------------GVAKLNIFFKELNYKSNSES
Human α                 ------------NYTVNNK----RN--------------------GVAKVNIFFKELNYKTNSES
Chicken α               ------------KYNITSK----RN--------------------GVAKVNIFFEEWNYKTNGES
Turtle α                ------------KYNITSK----RN--------------------GVAKVNIFFEEWKYKTNGES
Cow δ                   SSASIKSWPLPLPSSPSRA----RTEGPTSRSGAQPLSPEPCPSISLAKVNIFYQELNYRTVDET
Human δ                 -----------LPHQSHRQ----RS--------------------SLAKINIVYQELNYRSVEEA
Coelacanth δ            ------------GKNLTNN----RR--------------------DVSKLNIFFQKLSYESFDET
Chicken δ               ------------GYNSTSN----RK--------------------DIAKVTIYYKQLNYQSVNES
Turtle δ                ------------GYNSTHN----RR--------------------DIAKVNIFYQQLNYQSVDES
S. Lamprey β            ------------KHNVSRI-FNRKQ--------------------DIIKLNLFFQEFNSMTISES
J. Lamprey β            ------------KHNVSRQ--------------------------DIIKVNLFFQEFNSMTTSES
E. Shark β              ----------DSSPKVTVN----RN--------------------SILKLNMYFKENNFRTISES
Coelacanth β            ----------DLAINRTMN----RN--------------------GALKLNLYFQEFNYRTISES
Ropefish β              ----------DFSSNVTIN----RD--------------------GVMKMNLYFKEINYRSITES
W. Lungfish β           ----------DNSSDITMN----SD--------------------GVLRLNFYFNEFNYRVISES
A. Lungfish β           ----------DDSTNITMK----RD--------------------GVLKLNLYFKEFNYRVITES
Frog β                  ----------DSSHNITVN----RN--------------------GIVRLNIYFQEFNYRSISES
Cow β                   ----------DQSSNITLS----RK--------------------GIVKLNIYFQEFNYRTIEES
Human β                 ----------DQSTNITLS----RK--------------------GIVKLNIYFQEFNYRTIEES
Chicken β               ----------DMSTNVTLD----RN--------------------GIIKLNIYFQEYNYRTISES
Turtle β                ----------DLSTNVTLD----RN--------------------GIIKLNIYFQEYNYRTISES
S. Lamprey γ            ----------GLPANDTLK----PS--------------------DIAIVNIYFKDLTQKTISES
J. Lamprey γ            ----------GLPANDTLK----PS--------------------DIAIVNIYFKDLNQKTISES
Ropefish γ              ----------VLKKKQNIS----KE--------------------NFAKFSIFYKDLNLKTITES
E. Shark γ              ----------GRDGNKTLS----TS--------------------ELASLDIYYADLSVRNITEK
W. Lungfish γ           ------------GLNGTLS----KN--------------------DLVNLAIFYQDLNWRSLSES
A. Lungfish γ           ------------GLNNTLN----KN--------------------DLASIAIFYQDLNLRSLSES
Coelacanth γ            ----------SLKGNKGLS----KN--------------------DLVNLGIFYKDLNLRSISES
Frog γ                  ----------GEKLNKNLT----KN--------------------DLANLNIFYQDLNSRSISES
Cow γ                   ----------SQQIKKKLN----KT--------------------DLAKLLIFYKDLNQRSIMEN
Human γ                 ----------GRQVNKKLN----KT--------------------DLAKLLIFYKDLNQRSIMES
Chicken γ               ----------GQKINKKLN----KT--------------------DLANLMVFYKDLNERFISEN
Turtle γ                ----------GQKINKMLN----KT--------------------DLANLVVFYKDLNERFISEN
```

| | |
|---|---|
| Spotted Gar γ-like | PSFQVIDLVSSIGGLVGLWIGVSICTVAEFVE-LFLKVIIFIIKRVIR----------------- |
| Asian Arowana γ-like | PFITDIDLFSSVGGLVGLWVGVSLCTLAEFLE-FAVNVVLIVARGCLS----------------- |
| Coelacanth ASIC1 | KAYEVAGLLGDIGGQMGLFIGASILTILEILD-YLYEVFKDQVLGYFK----RKKRPKRSH---- |
| Catfish ASIC1 | KAYEVAGLLGDIGGQMGLFIGASVLTILEIFD-YLYEVFKDKVLGYFLR-KGRPRRSASDN---- |
| E. Lamprey ASIC1 | RAYEVAGLLGDIGGQMGLFIGASILTILEIFD-YLYEIIKYRILYYFR----RNKKQRNIS---- |
| E. Shark ASIC1 | KAYEVAGLLGDIGGQMGLFIGASLLTILELFD-YVYEVLKHKMCGVLR--MGKQQKRNNND---- |
| J. Medaka ASIC1 | KAYELAGLLGDIGGQMGLFIGASILTVLELFD-YLYEILKYKLCRCMK---KKHKSRNNND---- |
| Black Rock Cod ASIC1 | KAYELAGLLGDIGGQMGLFIGASLLTILELFD-YLYEVIKYKLCRCVK---KKHKGRNNND---- |
| Lancelet γ-like | PAYEVENLLGDLGGQLGLWVGMSCLSAMELLE-FLVDIAIIL------------WKKMSNR---- |
| Lancelet α-like | PAYRVENLLGDLGGQLGLWVGVSVMTILEVLE-LIVDVTQILLS------KARGKKKTRV----- |
| E. Shark α | SAISVMELLSSLGGEVSVWFGGSVLSGFELLE-LLLDFLVLSIALL----KASVWRGH------- |
| Frog δ | PTMSVNVLLSSMGGLWSWWFGSSVLSVAEIAE-LVLDTAAMVTIVIYQ--WKKQRANRNDG---- |
| Coelacanth α | PSINAATLLSNLGSQWSFWFGSSVLSVIEMVE-LLIDFFVLSTILLFR--HYCIQREENQD---- |
| S. Lamprey α | AATNVIWLLSNIGSQWSLWFGSSVLSWLEVGE-LGIDCCIMVFVLAYR--RRRSRAERRRA---- |
| J. Lamprey α | AATNVIWLLSNIGSQWSLWFGSSVLSWLEVGE-LGIDCCIMVFVLAYR--RRRSCAERRRARGTE |
| W. Lungfish α | PSFNVVTLLSNMGSQWSLWFGSSVLSVVEMGE-LVIDLIAVGIIVLRR----RQREKTKAT---- |
| A. Lungfish α | PSFTVVTLLSNMGSQWSLWFGSSVLSVVEMGE-LVFDLIAVGVIVLRR----RRREKCQAS---- |
| Ropefish α | PAFTAVMLLSNLGSQWSLWFGSSVMSVVELAE-LVFDLVAITLIFSAQ--KYFQWKNTESKVNTD |
| Frog α | PTINMAMLLSLLGSQWSLWFGSSVLSVVEMLE-LVIDFVIIGVMILLHRYYYKKANEGEET---- |
| Salamander α | PAINMVVLLSLLGSQWSLWFGSSVLSVVEMAE-LLFDVAAITVILYLQ--RRRKRQMDSVE---- |
| Cow α | PSVTMVTLLSNLGSQWSLWFGSSVLSVVEMAE-LIIDLLVITFLMLLRRFRSRYWSPGRGG---- |
| Human α | PSVTMVTLLSNLGSQWSLWFGSSVLSVVEMAE-LVFDLLVIMFLMLLRRFRSRYWSPGRGG---- |
| Chicken α | PAFTVVTLLSQLGNQWSLWFGSSVLSVMELAE-LILDFTVITFILAFR--WFRSKQ--------- |
| Turtle α | PAFTVVTLLSQLGNQWSLWFGSSVLSVVELAE-LILDFIAITFILSFR--WLRSRQ--------- |
| Cow δ | PVYSVPQLLSAMGSLWSLWFGSSVLSVVEVLE-LLLDAIALTLLLCCR--WLCGSRGQPRA---- |
| Human δ | PVYSVPQLLSAMGSLCSLWFGASVLSLLELLE-LLLDASALTLVLGGR--RLRRAWFSWPR---- |
| Coelacanth δ | PSISAVTILSQMGNLWSFWFGSSVLSVIELIE-LILDVIAMSFILTFK--WHKLKK--------- |
| Chicken δ | PLLSDNLLLSSMGSQWSLWFGSSVLSVVEMLE-LLIDTLVLSLLFCYQ--RFRSKTLNVAR---- |
| Turtle δ | PVYTVNLLLSNMGSQWSLWFGSSVLSVVEMFE-LLLDIMVLSLIFCYR--RFKAKKTLKMA---- |
| S. Lamprey β | PAQTIVTLLSNLGGQFGFWMGGSVLCIIEFIE-IIIDCVWIGMIKASN--DVRERRKTSRK---- |
| J. Lamprey β | PAQTIVTLLSNLGGQFGFWMGGSVLCIIEFIE-IIIDCVWIGIIKASH--DVRERRKTSRK---- |
| E. Shark β | KAQTLVWLLSNLGGQFGFWMGGSVLCIVELLE-VMLDCVWIMAIRGAR--LYMGHRRRRSL---- |
| Coelacanth β | AATDISWLVSNLEGQFGFWMGGSILCIIEFLE-IIIDCVWITIIKLVI--WYRDRKHKKAQ---- |
| Ropefish β | ASTTVVWLLSNLGGQFGLWMGGSVICIVEFGE-IIIDCLWITIIKLIM--WNRNWKLKKAQ---- |
| W. Lungfish β | AATNVVWLLSNLGGQFGFWMGGSVLCIIEFGE-IIIDCIWIAIIRFVI--WQKNRKKMQLP---- |
| A. Lungfish β | VATNVVWLLSNLGGQFGFWMGGSVLCIIEFGE-VFIDCIWIAVIRFVK--WYKNRKERQVQ---- |
| Frog β | EATNVVWLLSNLGGQFGFWMGGSVLCIIEFGE-IIIDCMWITILKFLA--WSRNRRQRRKR---- |
| Cow β | AANNIVWLLSNLGGQFGFWMGGSVLCLIEFGE-IIIDFVWITIIKLVA--LAKSVRQKRAQ---- |
| Human β | AANNIVWLLSNLGGQFGFWMGGSVLCLIEFGE-IIIDFVWITIIKLVA--LAKSLRQRRAQ---- |
| Chicken β | AATTIVWLLSSLGGQFGFWMGGSVLCLIEFGE-IIIDSLWITVINIIS--WCKGLKQKRVR---- |
| Turtle β | AATTIVWLLSSLGGQFGFWMGGSVLCLIEFGE-IIIDFLWITIINIIS--WCKGLKQKRAR---- |
| S. Lamprey γ | PASSIVTLLSNLGGLLGLWLSCSMLCVVEVLEIFCVDFPWILLKKLLT--TCSSAFASLIR---- |
| J. Lamprey γ | PASSIVTLLSNLGGLLGLWLSCSMLCVVEVLEIFCVDFPWILLKKLLA--TCRRTFASLVR---- |
| Ropefish γ | PLNNIVTLLSNVGGQLGLWLSCSIVCVIEIIEVFFLDAPWILVRQIIR--SCQIRCRERQQ---- |
| E. Shark γ | PANTMVTLLSNFGGQLGLWMSCSVICVIEIVEVLLVDALWVLVRSAGQ--RVRRWW--------- |
| W. Lungfish γ | PANSIVTLLSNFGGQNGLWMSCSVVCFIEIWEVFLVDILTILVRNWFR--KAKLWWNKKKE---- |
| A. Lungfish γ | PANSIATLLSNMGGQLGLWMSCSIVCFLEMWEVFLVDILTIIARYWLH--RGRQWWRKRKE---- |
| Coelacanth γ | PANNIVTLLSNFGGQLGLWLSCSVVCVLEIIEVFFIDAFWIVLRQTTQ--KARDWWTKRK----- |
| Frog γ | PTYNIVTLLSNFGGQLGLWMSCSMICVLEIIEVFFIDSFWVVLRQRWR-----NWWENRKE---- |
| Cow γ | PANSIEQLLSNIGGQLGLWMSCSVVCVIEIIEVFFIDSLSIIARHQWH--KAKGWW----A---- |
| Human γ | PANSIEMLLSNFGGQLGLWMSCSVVCVIEIIEVFFIDFFSIIARRQWQ--KAKEWW----A---- |
| Chicken γ | PANTLVILLSNFGGQLGLWMSCSVVCVIEIIEVFFIDSFSIVMRRQWQ--KAKKWWNHRKR---- |
| Turtle γ | PANNIVILLSNFGGQLGLWMSCSVVCVIEIIEVFFIDSFFIVTRRRWQ--KAKKWWRDRKA---- |

| | |
|---|---|
| Spotted Gar γ-like | -KNKEAPLNPYM-----------------------------IAHQTFKSSV-------------- |
| Asian Arowana γ-like | -SRSQGPASSAL----------------------------------------------------- |
| Coelacanth ASIC1 | -SDNLSTCDTLR---------------------------------SHSDSLGFTPNMLPR----- |
| Catfish ASIC1 | -LDYSENPTSPG-----------------------------------VTPNHTPRAHATH----- |
| E. Lamprey ASIC1 | -DNSVPMTSSYA---APQGHTQAAIH--------------------------------------- |
| E. Shark ASIC1 | -KGVTLSLDDIK--RHNPC----------------------ESLRGHPTGMSYTANMLPH----- |
| J. Medaka ASIC1 | -RGAVLSLDDVK--RHAPCE-------------------------NLRTPSTYPGNMLPH----- |
| Black Rock Cod ASIC1 | -RGAVLSLDDVK--HHDPCD-------------------------NLRTPSTYPGNMLPH----- |
| Lancelet γ-like | -EKTTSRNNVVN--VAETG----------------------KIGTSNGHTIPIHMTEVM------ |
| Lancelet α-like | --------------IDISGHM-------------------------------------------- |
| E. Shark α | ----------------------------------------------------------------- |
| Frog δ | -MNGSNAASNVC-----------------------------SVPYIDSKTFSTPQTGLSE----- |
| Coelacanth α | -PEITISTVSYP--HYANE----------------------NSEFNSEESMGNHFDVVAD----- |
| S. Lamprey α | -RGTGDSEAAPP----------------------------------------------------- |
| J. Lamprey α | DSEAAPPPPPSF--REALGCANAAYDGANDGADDG------HDGDGGGVAIGGPTERPPPWRRGS |
| W. Lungfish α | -DDTEDNSPSET--YVHPRQENS------------------DNEHRERAPNRIEVVAEIS----- |
| A. Lungfish α | -SDGEGTSDSTA--GTHRGQ---------------------ENASRSGRDVACNRFVVVA----- |
| Ropefish α | GHQEPNTHNAVSNENHQEFDSLGVNFAFEAEDATHSIQSIPSDSPNVEMTTSFQFDVVAD----- |
| Frog α | -TVVPTPAPAFA--DLEQ-----------------------QVPHIPRGDLSQRQISVVA----- |
| Salamander α | -EDSQAPTPTLP---RFEGHGNPEFQ------------------SEEEPSHQFRVVADIT----- |
| Cow α | -KGTQEVASTPA--ASLPSS---------------------FCPHPAFFSSSPPDPAISP----- |
| Human α | -RGAQEVASTLA--SSPPSH---------------------FCPHPMSLSLSQPGPAPSP----- |
| Chicken α | -WHSSPAPPPNS-------HDNTAFQ---------------DEASGLDAPHRFTVEAVVT----- |
| Turtle α | -LLASAVPPPGA-------HDNTAFQ---------------PEPSGPSAPHRFTVEAVVT----- |
| Cow δ | -ATRVHPPSQRP------------------------------------ASGPVAADTTSN----- |
| Human δ | -ASPASGASSIK--PEAS-----------------------QMPPPAGGTSDDPEPSGPH----- |
| Coelacanth δ | ----------------------------------------------------------------- |
| Chicken δ | -TPSIPSVSLTL--ESYRVVQEAGN----------------GTAPAHGHTSGVPMAVANS----- |
| Turtle δ | -QPLAISSVTLTLENYRAVHEDLAADWNSIWTNQN------VGTVAKTKNGDLPFHSICN----- |
| S. Lamprey β | -PRYSDEPPTLS--SIVQGQGNSGFEMEERGPPG-------EAPEANGSAAQPPAAAAEQ----- |
| J. Lamprey β | -PRYSDEPPTLS--SIVQGQGNSGFEMEEREPPR-------EAPDANGSAAQPPTAAAEQ----- |
| E. Shark β | -ARHPPPVPTVA---------------------------------QCVEEQEHRAQSERP----- |
| Coelacanth β | -AQYSGPPPSVS--QLARAHTDTGFQ------HD-------STDINYGTEAYCNEAYIPP----- |
| Ropefish β | -SQYMGPPPSIS--QLAEGHMNTGFE------PD-------NVTSDSGQSLQTMSSCEHP----- |
| W. Lungfish β | -QLYNDPPPTVS--ELVEGISNQGFQ---------------PDIINSCTSQPQPPDLHIP----- |
| A. Lungfish β | -AQYADPPPTVS--ELVEGYTNQGFQ---------------PDIINSCSPQAQPPDLYLP----- |
| Frog β | -PQYSDPPPTVS--ELVEAHTNSGFQ--------------------HDDGDHVPVD--------- |
| Cow β | -ARYEGPPPTVA--ELVEAHTNFGFQ------PD-------LATPGPDVE-AYPHEQNPP----- |
| Human β | -ASYAGPPPTVA--ELVEAHTNFGFQ------PD-------TAPRSPNTG-PYPSEQALP----- |
| Chicken β | -ARYPDTPPTVS--ELVEAHTNLGFQ-------------------HEEAGTETQGEALPP----- |
| Turtle β | -AQYPDTPPTVS--ELVEAHTNLGFQ------HE-------DTNITPCDE-VPPPGALPP----- |
| S. Lamprey γ | -GPDPAPSVHFP--VQLPVGGSPA-----------------EDPPTFHTAMQCPREPIPM----- |
| J. Lamprey γ | -GPDPAPSVHFP--VQLPVGGSPA-----------------EDPPTFHTAMQCPREPIPT----- |
| Ropefish γ | ----RPTLPTVS--GGIPVD---------------------EDPPTFNSALRLPQPNLLE----- |
| E. Shark γ | ---QRPEQEEVP--AHGTAHGITVYL------ED-------EDPPTFNAALRLP--RQCP----- |
| W. Lungfish γ | -GQ-EQVYN--Q--NRHTGHDNPVCV------ED-------EDPPTFHTAMQLPCVQTGE----- |
| A. Lungfish γ | -RQMQQPSP--P--DHDTGHHNPVCI------DD-------EDPPTFHTAMQLPCVQTGP----- |
| Coelacanth γ | -VRQAQPSLAAP--TDYAGQHNPVYVSD-------------EDPPTFSTAVHLPHSESCP----- |
| Frog γ | -NQAEDTPEIPV--PTMTGHDNPLCVDNPICLGE-------EDPPTFNSALQLPQSQDSH----- |
| Cow γ | -RRRAPACPEAP--RAPQGRDNPSLD------ID-------DDLPTFTSALSLPPAPGSQ----- |
| Human γ | -WKQAPPCPEAP--RSPQGQDNPALD------ID-------DDLPTFNSALHLPPALGTQ----- |
| Chicken γ | -DETGKPPEVGD--AEQQGHDNPAC-------SD-------EDLPTFNTALRLPLPQEGH----- |
| Turtle γ | -APAQVNAPAKE------GHDNPVCID--------------EDLPTFNTALHLPLPQENH----- |

```
Spotted Gar γ-like       -----------------------------------------------------------------
Asian Arowana γ-like     ----------------------------------MAGREGPPS----------------------
Coelacanth ASIC1         -------------------------------------HPPLGNFEEFAC----------------
Catfish ASIC1            ------------------------------------SGVTRTASDSRRTCYLVTRL---------
E. Lamprey ASIC1         ------------------------------------PHGAHAKYEDFTC----------------
E. Shark ASIC1           -------------------------------------HPPRNTFEDFTC----------------
J. Medaka ASIC1          -------------------------------------HPGQGNFEDFTC----------------
Black Rock Cod ASIC1     -------------------------------------HPGQANFEDFTC----------------
Lancelet γ-like          -----------------------------------------------------------------
Lancelet α-like          -----------------------------------------------------------------
E. Shark α               -----------------------------------------------------------------
Frog δ                   -----------------------------------------KELESKTNDPKFNGEINTFS----
Coelacanth α             -------------------------------------VSAPPAYETLDLDLPPSAAQCITGCKCV
S. Lamprey α             ---------------------------------------PPPSF---------------------
J. Lamprey α             LTRNACSFCVVADISCPPAAAAKPVVGASREDAAAAPEALPPDYGSLRRVPEGYVAVSSDPGGPP
W. Lungfish α            ---------------------------------------PPPAYDSLELDTPVACSADCSCTRRM
A. Lungfish α            ------------------------------------EISPPPAYDTLQLDVPVACAPDCECTQHV
Ropefish α               -------------------------------------ISPPPAYDSLNLECSSLSQTIKGCNAEC
Frog α                   ------------------------------------DITPPPAYESLDLRSVGTLSSRSSSMRSN
Salamander α             ---------------------------------------PPPAYDSLELHAREECDDSCSCSHRS
Cow α                    -----------------------------------ALSAPPPAYATLGPHPAPSGLAEASTSAHA
Human α                  -----------------------------------ALTAPPPAYATLGPRPSPGGSAGASSSTCP
Chicken α                ---------------------------------------TLPSYNSLEPCGPSKDGETGLE----
Turtle α                 ---------------------------------------TLPSYNSLELCGQNRDAEIGVE----
Cow δ                    --------------------------------------APGPGCLHLPRCCRDFSRSLG------
Human δ                  -----------------------------------LPRVMLPGVLAGVSAEESWAGPQPLETLDT
Coelacanth δ             -----------------------------------------------------------------
Chicken δ                -------------------------------------SDPHPAQLSSKAIPEHCPDVVLNGFRYM
Turtle δ                 -------------------------------------KTPTPELNPDVVLNGFRHIKDHSIEIDL
S. Lamprey β             --------------------------------QPDVPGTPPPHYDTLRISKTELHDEINSDDDGE
J. Lamprey β             ---------------------------------QDVPGTPPPHYDTLRISKTELHDEINSDDDSE
E. Shark β               -----------------------------------VPSTPPPRYDSLHICSLTEPGPPEAGGKIV
Coelacanth β             --------------------------------REPTPGTPPPNYDSLRVQPVENTEQISDSEEN-
Ropefish β               --------------------------------VHPIPGTPPPHYDSLRLKSVRVNNE--------
W. Lungfish β            -------------------------------TTLEVPGTPPPHYDSLRIQPIDMEQQSDNEDF--
A. Lungfish β            -------------------------------TTLEIPGTPPPKYDSLRVHPIDTEHHSDSEDL--
Frog β                   -----------------------------------IPGTPPPNYDSLRVNTAEPVSSDEEN----
Cow β                    -----------------------------------IPGTPPPNYDSLRLQPLDVIESDSEGDAI-
Human β                  -----------------------------------IPGTPPPNYDSLRLQPLDVIESDSEGDAI-
Chicken β                -----------------------------------EPGTPPPNYDSLRVQPSHNPGTDSDIECEE
Turtle β                 -----------------------------------EPGTPPPNYDSLRVDPRAGIDSDSDAEAS-
S. Lamprey γ             ------------------------------------PNTPPPQYNTLRLRQIAGYVPDGGSDGE-
J. Lamprey γ             ------------------------------------PNTPPPQYDTLRLRQIAGYVPDEGSDGED
Ropefish γ               -----------------------------------VPKTPPPNYNTLRIHNVFSHMNDEDEHDTI
E. Shark γ               ------------------------------------PTAPPPNYEMLQQCHGFNGGPEEERF---
W. Lungfish γ            -----------------------------------VPSTPPPQYDALRIQNVFDEQFSDTEVN--
A. Lungfish γ            -----------------------------------VPSTPPPQYNALRIQSVFDEQVSDTEVN--
Coelacanth γ             -----------------------------------VPKTPPPTYDALRIQTAFAEQISDTEDNEY
Frog γ                   -----------------------------------VPRTPPPKYNTLRIQSAFQLETIDSDEDVE
Cow γ                    -----------------------------------VPGTPPPRYNTLRLERAFSSQLTDTQTTFP
Human γ                  -----------------------------------VPGTPPPKYNTLRLERAFSNQLTDTQMLDE
Chicken γ                -----------------------------------PPRTPPPNYSTLRLETAFTEQLPDTLEAGQ
Turtle γ                 -----------------------------------VPRTPPPNYSTLKLDAAFTDPLPDTLEGSC
```

| | |
|---|---|
| Spotted Gar γ-like | ----------------------------------------- |
| Asian Arowana γ-like | ----------------------------------------- |
| Coelacanth ASIC1 | ----------------------------------------- |
| Catfish ASIC1 | ----------------------------------------- |
| E. Lamprey ASIC1 | ----------------------------------------- |
| E. Shark ASIC1 | ----------------------------------------- |
| J. Medaka ASIC1 | ----------------------------------------- |
| Black Rock Cod ASIC1 | ----------------------------------------- |
| Lancelet γ-like | ----------------------------------------- |
| Lancelet α-like | ----------------------------------------- |
| E. Shark α | ----------------------------------------- |
| Frog δ | ----------------------------------------- |
| Coelacanth α | HCASFISHEVEDLMSDLAGDP-------------------- |
| S. Lamprey α | ----------------------------------------- |
| J. Lamprey α | RRGAGAPLLSPRVSALAASGKEVLRRRSASLNVVSFAIEEA |
| W. Lungfish α | SQTSIKSHTSNSSNTEESTSEGPTAL--------------- |
| A. Lungfish α | SHASVHSQAPCSSQPEQEASEGPTVL--------------- |
| Ropefish α | QCSRRLCEINEKD---------------------------- |
| Frog α | RSYYEENGGRRN----------------------------- |
| Salamander α | SIRSNMSVHSRASSAATNA---------------------- |
| Cow α | PGEP------------------------------------- |
| Human α | LGGP------------------------------------- |
| Chicken α | ----------------------------------------- |
| Turtle α | ----------------------------------------- |
| Cow δ | ----------------------------------------- |
| Human δ | ----------------------------------------- |
| Coelacanth δ | ----------------------------------------- |
| Chicken δ | KDSSLGGEINH------------------------------ |
| Turtle δ | NS--------------------------------------- |
| S. Lamprey β | FV--------------------------------------- |
| J. Lamprey β | FVS-------------------------------------- |
| E. Shark β | VEVGGEEAGEEWGEEGCRL---------------------- |
| Coelacanth β | ----------------------------------------- |
| Ropefish β | ----------------------------------------- |
| W. Lungfish β | ----------------------------------------- |
| A. Lungfish β | ----------------------------------------- |
| Frog β | ----------------------------------------- |
| Cow β | ----------------------------------------- |
| Human β | ----------------------------------------- |
| Chicken β | QRPAANHHGDASVWAE------------------------- |
| Turtle β | ----------------------------------------- |
| S. Lamprey γ | ----------------------------------------- |
| J. Lamprey γ | ----------------------------------------- |
| Ropefish γ | TPIDDSIVHDNTMTINQRRRKENPFLII------------- |
| E. Shark γ | ----------------------------------------- |
| W. Lungfish γ | ----------------------------------------- |
| A. Lungfish γ | ----------------------------------------- |
| Coelacanth γ | ----------------------------------------- |
| Frog γ | RL--------------------------------------- |
| Cow γ | H---------------------------------------- |
| Human γ | L---------------------------------------- |
| Chicken γ | H---------------------------------------- |
| Turtle γ | H---------------------------------------- |
